# Aldehyde dehydrogenase 1A3 detection in extracellular vesicles from breast cancer cell lines by nano-flow cytometry

**DOI:** 10.64898/2026.06.17.732934

**Authors:** A Pallarès-Rusiñol, R Pequerul, L Costa-Sastre, M Tuxans, A Constantinescu, M Pérez-Alea, M.I Pividori, J Farrés, M Martí

**Author notes:** Corresponding authors. E-mail addresses (J. Farrés), (M. Martí).

## Abstract

Exosomes are nanosized extracellular vesicles that carry bioactive molecules reflective of their cells of origin. Developing methods to detect functional enzymatic activity within exosomes can provide a new generation of rapid and informative diagnostic tools. Aldehyde dehydrogenase (ALDH) enzymes, particularly ALDH1A3, are overexpressed in several cancers and contribute to tumor aggressiveness and drug resistance. However, their presence and functionality in cancer-derived exosomes remain poorly characterized. Here, we developed a nano–flow cytometry–based method to detect ALDH activity directly within individual exosomes derived from breast cancer cell lines (SKBR3, MDA-MB-231, and MCF7). Exosomes were isolated by differential ultracentrifugation and validated by nanoparticle tracking analysis, cryogenic transmission electron microscopy, and bead-based immunophenotyping of canonical markers. ALDH enzymatic activity was detected using a resorufin-based fluorescent substrate capable of crossing the exosomal membrane. To ensure specificity, assays were performed in the presence or absence of a selective ALDH inhibitor, confirming that the fluorescent signal originated from ALDH activity within the vesicles. This work provides the first functional evidence of ALDH1A3 enzymatic activity in cancer-derived exosomes and establishes a proof-of-concept platform for rapid, activity-based detection of exosomal enzymes, opening new perspectives for exosome-based diagnostics in breast cancer.

## 2. Introduction

Aldehyde dehydrogenases (ALDHs) constitute a superfamily of NAD(P)^+^-dependent enzymes that catalyze the irreversible oxidation of diverse endogenous and exogenous aldehydes to their corresponding carboxylic acids. In addition to this canonical function, several ALDH family members also display cofactor-independent esterase activity, reflecting the broad catalytic versatility of this enzyme family. (Zhou et al., 2019).

The human genome encodes 19 ALDH isoforms, grouped into distinct families and subfamilies based on amino acid sequence identity, with functional diversity arising from differences in tissue distribution, subcellular localization, and substrate specificity. Among them, the ALDH1A subfamily (ALDH1A1, ALDH1A2 and ALDH1A3) comprises cytosolic tetrameric enzymes that primarily convert retinaldehyde to retinoic acid and also oxidizes aliphatic aldehydes from lipid peroxidation and xenobiotic-derived aldehydes formed during drug metabolism. These isoforms share approximately 70% amino acid identity and overlapping substrate preferences, making their differential functional characterization and the development of selective inhibitors particularly challenging (Pequerul et al., 2020). Despite their structural similarity, ALDH1A isoforms exhibit distinct physiological roles related to oxidative stress response, lipid hormone metabolism, retinoic-acid-dependent cell differentiation, immunomodulation, and metabolic adaptation. These functions in normal tissues can be co-opted in malignant contexts, where they contribute to tumor survival, proliferation and therapeutic resistance. Accordingly, elevated ALDH1A activity has been widely associated with poor clinical outcomes and is routinely used as a hallmark of cancer stem cells (CSCs), reflecting their persistence and chemoresistance (Zhou et al., 2019).

The oncogenic contribution of ALDH enzymes, and particularly ALDH1A subfamily, has been increasingly investigated over the past decade. Their involvement extends beyond CSC maintenance to include direct detoxification of cytotoxic aldehydes, immunomodulation and metabolic reprogramming favoring glycolysis over oxidative phosphorylation (Hirpara et al., 2024; Kang et al., 2016).

Among ALDH1A isoforms, ALDH1A3 predominates in breast cancer, particularly in basal-like and triple-negative subtypes. Its expression correlates with aggressive phenotypes, therapy resistance and poor prognosis (Marcato et al., 2015). ALDH1A3 also supports the stem-like and invasive features of circulating tumor cells (CTCs), enhancing their adhesion, migration and metastatic colonization (McLean et al., 2023; Yamashita et al., 2020). Recently, we showed that ALDH1A3 expression stratifies breast cancer subtypes and associates with advanced stage and lymph node positivity, reinforcing its link to metastatic progression and therapeutic resistance (Pequerul et al., 2025). These observations support ALDH1A3 as a biomarker and therapeutic target in aggressive disease.

Extracellular vesicles (EVs) mediate local and systemic cell-to-cell communication, shaping the tumor microenvironment and conditioning distant tissues, for example pre-metastatic niches. Among EVs, exosomes are a distinct, nanoscale population (30-200 nm) generated by the endosomal pathway and released by most cell types (Gurung et al., 2021; Johnstone et al., 1987). By encapsulating nucleic acids, lipids, and proteins reflective of their cells of origin, exosomes act as vehicles of functional transfer and reservoirs of disease-relevant biomarkers. Notably, exosomal cargo frequently includes glycolytic enzymes and other metabolic mediators that can reprogram the energy balance of recipient cells and remodel the microenvironment toward pro-survival and immune-evasive states, thereby promoting tumor progression, persistence and therapy resistance (Fridman et al., 2022; Keerthikumar et al., 2016; Vahabi et al., 2023; Wang et al., 2020a; Xu et al., 2023; Yang et al., 2020) Consistent with this, recent studies implicates exosomes as active mediators of tumor immune evasion (Saadh et al., 2025; Whiteside, 2025; Zhang et al., 2025).

Various ALDH isoforms have been detected in extracellular vesicles derived from multiple pathological contexts, including ALDH1A1 in colon, pancreatic, breast, and lung cancers (Emmink et al., 2013; Gonzalez-Callejo et al., 2023; He et al., 2024; Hirpara et al., 2024; Servage et al., 2020; Shang et al., 2025); ALDH1A3 in prostatic secretions (Principe et al., 2013); ALDH2 in nasopharyngeal carcinoma (Yan et al., 2024); ALDH3A1 in lung cancer (Wang et al., 2020b); and ALDH7A1 in oral squamous cell carcinoma and their paired lymph node metastatic cells (Busso-Lopes et al., 2021). However, in most reports, these identifications relied on protein-level detection methods (e.g., immunoblotting), providing limited information on whether the enzyme molecules retained catalytic functionality. For metabolic enzymes such as ALDHs, functional activity, rather than mere presence, is essential to infer biological relevance.

To date, ALDH1A3 enzymatic activity has not been demonstrated within the exosomal cargo. The present study aimed to bridge this gap by integrating two complementary strategies: 1) Enzymatic activity-based detection to functionally infer the presence of exosomal ALDH1A3, and 2) Tracer-labelled exosomes analysis by nano-flow cytometry for single-particle assessment.

We isolated and characterized exosomes from breast cancer cell lines (SKBR3, MDA-MB-231 and MCF7) using differential ultracentrifugation and validated their integrity by nanoparticle tracking analysis, cryo-electron microscopy and bead-based immunophenotyping. Exosomal ALDH1A3 activity was then evaluated using the fluorescent substrate resorufin propionate (RP), capable of permeating exosome membranes and reacting with intravesicular ALDH. Because RP can also be processed by esterase enzymes, we further applied a specific ALDH inhibitor to distinguish ALDH-dependent fluorescence, enabling the first demonstration of functional ALDH1A3 activity within cancer-derived exosomes.

## 3. Materials and Methods

### 3.1 Instrumentation

Nanoparticle tracking analysis (NTA) was performed using the NanoSight LM10-HS system with a tuned 405 nm laser (NanoSight Ltd, Malvern, GB). Cryogenic transmission electron microscopy images were acquired by a Jeol JEM 2011 (JEOL USA Inc, US). Spectrophotometric measurements were performed on a Tecan Infinite m200 PRO (Tecan Group Ltd, Männeford, CH) microplate reader controlled by Magellan v7.0 software. Conventional and nano-Flow cytometry measurements were performed using a Cytoflex LX (Beckman Coulter Inc, Indianapolis, IN, US) and analyzed with the integrated software Cytexpert and FlowJo analysis software (FlowJo LLC, NJ, US).

### 3.2 Chemical reagents

Tosylactivated magnetic particles (MPs) (Dynabeads M450 Tosylactivated, ref. 14013), FITC-labelled mouse monoclonal antibodies against tetraspanins, antiCD9 (ref. MA119557), antiCD63 (ref. MA119602), antiCD81 (ref. A15753), as well as mouse monoclonal antiVinculin (ref.MA5-11690) and PE-labelled goat anti-Rabbit secondary antibody (antirabbit-PE, ref. P2771MP) were purchased from Thermo Fisher Scientific (Waltham, MA, US). Rabbit polyclonal antibody against ALDH1A3 (ref. GTX110784) was purchased from Genetex (Irvine, CA, US). Cy5-labelled goat anti-Mouse (antimouse-Cy5, ref. ab97037) was purchased from Abcam (Cambridge, GB). For exosome membrane staining, Cell Trace CFSE green kit (ref. C34554) and Cell Trace Violet kit (ref.C34557) were purchased from Thermo Fisher Scientific. For the ELISA quantification, HRP-conjugated antiCD63 antibody (ref. NBP2-42225H) was purchased from Novus Biologicals (Bio-Techne R&D Systems SLU, Madrid, ES), and Pierce TMB substrate kit (ref. 23227) was purchased from Thermo Fisher Scientific. For the protein quantification, Pierce BCA Protein Assay kit (ref. 23227) was purchased from Thermo Fisher Scientific. For cellular lysis and protein extraction, M-PER reagent (ref. 78501) was also purchased from Thermo Fisher Scientific. For the cell culture, Dulbecco’s Modified Eagle’s Glutamax (DMEM, ref. 31966-021) medium, and fetal bovine serum (FBS, ref. 26140079) were purchased from Gibco (Thermo Fisher Scientific). The composition of all buffers and solutions is described in S1 (Supp. Data).

For ALDH activity studies, hexanal (Sigma Aldrich, St. Louis, MO, USA, ref. 115606) and nicotinamide adenine dinucleotide in oxidized (NAD^+^; Apollo Scientific, Bredbury, UK, ref. BIB3014) and reduced (NADH; Apollo Scientific, ref. BIB3012) forms were used. RP and the selective ALDH1A3 inhibitor ABD0305 were synthesized in-house as previously described (Ceylan et al., 2022; Pequerul et al., 2025) and employed for the enzymatic activity as substrate and specific control, respectively (Figure 1).

**Figure 1.**
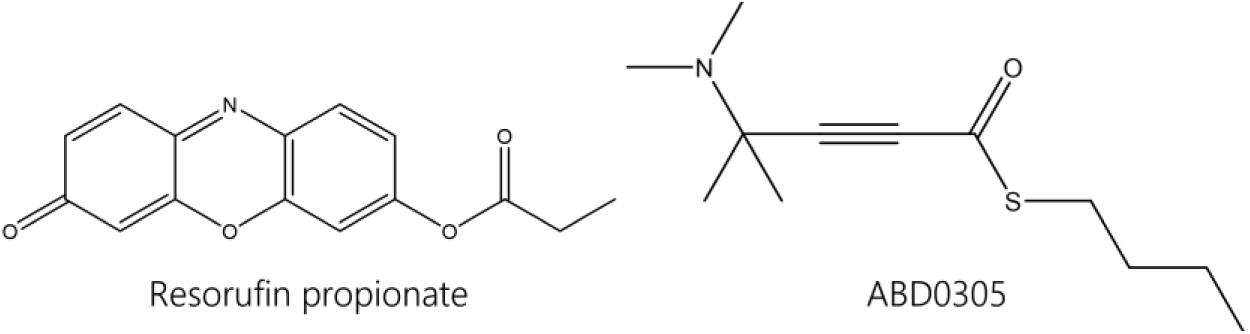
Molecular structures of resorufin propionate and inhibitor ABD0305.

### 3.3 Cell culture, exosome isolation and purification

Breast cancer cell lines SKBR3 (ATCC, ref. HTB-30), MDA-MB-231 (ATCC, ref. HTB-26) and MCF7 (ATCC, ref. HTB-22) were used. Expansion of cell population was carried out from 5 x 10^6^ cells in T-175 flask containing 35 mL of Dulbecco’s Modified Eagle’s medium. The media were supplemented with 10 % exosome-depleted fetal bovine serum (FBS) and 100 U mL^-1^ penicillin-streptomycin. The temperature was maintained at 37 °C in a humidified, concentrated CO_2_ (5 %) atmosphere. Once cells reached approximately 95 % confluence on the T-175 flask, the culture supernatant was removed and stored at –21°C until to exosome isolation. Exosomes were purified from cell culture supernatant by differential ultracentrifugation, as previously described by our group with minor modifications.(Moura et al., 2020). Full purification steps are provided in Supplementary Data S2. Pellets were resuspended in 25 mmol L^−1^ HEPES (pH 7.4, sterile-filtered through 0.22 µm membranes) and stored at –21°C.

### 3.4 Capillary electrophoresis-based immunodetection of ALDH isoforms in breast cancer cells

Cells were lysed in RIPA buffer containing phosphatase and protease inhibitors (Calbiochem, San Diego, CA, US). Total protein extracts were analyzed by capillary electrophoresis, using the WES Simple Western system according to manufacturer’s specifications (Bio-techne, San Jose, CA, USA). Immunodetection followed standard WES protocols, and chemiluminescence quantification and imaging were performed using Compass software for WES Simple western instruments (Bio-techne). Antibodies were: anti-ALDH1A2 (ref. 13951-1-AP), anti-ALDH1B1 (ref. 15560-1-AP), anti-ALDH2 (ref. 15310-1-AP) and anti-ALDH3B1 (ref. 15578-1-AP) from Proteintech (Martinsried, Planegg, GE); anti ALDH1A1 (ref. MAB5869) from R&D Systems; and anti-ALDH1A3 (ref. NBP2-15339) from Novus Biologicals.

### 3.5 Intracellular staining of ALDH in breast cancer cells

According to literature (Zanoni et al., 2022), ALDH enzymes localized to multiple cellular compartments (nucleus, cytoplasm, endoplasmic reticulum), with no evidence for plasma-membrane localization. To determine the intracellular presence of ALDH, cells were fixed with paraformaldehyde and stained in the presence of saponine to permeabilize membranes. A rabbit anti-ALDH1A3 primary antibody was used, followed by a PE-conjugated anti-rabbit secondary antibody. As an intracellular control, a mouse anti-vinculin antibody was used and detected with a Cy5-conjugated anti-mouse secondary antibody. Samples were analyzed by flow cytometry. Full experimental details of the assay are provided in Supplementary Data S3.

### 3.6 Fluorometric determination of ALDH1A3 enzymatic activity

Breast cancer cells and exosomes were lysed using M-PER™ extraction reagent, following the protocol provided by the manufacturer, and extracts were quantified by BCA protein assay kit. As standard quantification method for the ALDH activity in cells and exosomes, a fluorometric assay based on the oxidation of hexanal using NAD^+^ as cofactor was used (Pequerul et al., 2022). Enzymatic activity was monitored using a Cary Eclipse (Varian, Palo Alto, CA, US) fluorimeter at 25°C. All reactions were performed in quartz cuvettes in a final volume of 1 mL, using 50 mM HEPES (Merck, Darmstadt, GE), 50 mM of MgCl_2_, 5 mM DTT, pH 8.0, as the reaction buffer. Then, a 0.5 mM NAD^+^ cofactor (Apollo Scientific), and 250 μM hexanal substrate (Merck) were added to start the reaction. Fluorescence of NADH was followed at 460 nm with excitation at 340 nm using 10 nm for the excitation and emission spectral bandwidths. Five μM NADH was added to the reaction mixture as an internal standard to obtain absolute reaction rates. Specific activity was expressed in milliunits (mU) per mg of total protein of the lysate,1 mU defined as 1 nmol of product formed per min. To find the linear range of the enzymatic reaction, different dilutions of cells extract or exosomes were incubated. In the case of cell extracts, the fluorescence of the reaction was recorded for 100 min, with measurements each 30 s. Meanwhile in the case of exosomes, more time (4 hours) and higher temperature (37°C) were applied to increase the reaction kinetics. For statistical analysis, unpaired t tests with Welch’s correction were performed. Statistical significance was set at p values below 0.05.

### 3.7 Characterization of EVs by nanoparticle tracking analysis, cryogenic transmission electron microscopy and BCA protein assay

The size distribution and concentration of particles were estimated by NTA. The purified exosomes were diluted in filtered PBS buffer solution, between 100 and 500-fold depending on the sample initial concentration. NanoSight NTA software analyzed raw data videos by triplicate during 60 s with 25 frames/s. Cryogenic TEM images were collected by a Jeol JEM 2011 (JEOL USA Inc, US) transmission electron microscope at an accelerating voltage of 200 kV. Exosomes were maintained at -182 °C with liquid ethane during the whole process. The total protein concentration of the EVs was estimated using the Bicinchoninic acid protein assay (BCA), following the manufacturer instructions, using bovine serum albumin (BSA) standards in HEPES buffer solution. The spectrophotometric measurements were done at 562 nm. All experimental details are described in S4 (Supp. Data).

### 3.8 Characterization of exosomes by bead-based flow cytometry assay

Conventional flow cytometer was used to estimate the presence of general protein markers of exosomes in the surface of EVs derived from SKBR3, MDA-MB-231 and MCF7 breast cancer cell lines. Specifically, the expression of tetraspanin receptors CD9, CD63 and CD81 was determined, following MISEV2018 guidelines (Théry et al., 2018), to confirm the presence of exosomes The bead-based flow cytometry relies on the immobilization of exosomes on the surface of magnetic particles, to increase its size within the resolution of the flow cytometer. To achieve that, exosomes were covalently immobilized on MPs, followed by the direct labelling with FITC-modified antiCDX mouse monoclonal antibodies (being CDX either CD9, CD63 or CD81 biomarkers). All experimental details are described in S5 (Supp. Data).

### 3.9 Sandwich ELISA for the determination of ALDH

To determine the presence of ALDH1A3 on exosome membrane, exosomes derived from SKBR3 breast cancer cell line were subjected to sandwich ELISA. As a control for the exosome membrane staining, the antiCD81 antibody was added in the assay as general exosome membrane biomarker. Specific antibodies against ALDH1A3 and antiCD81 were used to capture the exosomes, further labelled by HRP-conjugated antiCD63 antibodies, and revealed with TMB and H_2_O_2_. The spectrophotometric measurements were done at 450 nm. All experimental details of the assay are provided in S6 (Supp. Data).

### 3.10 Nano-Flow cytometry studies of breast cancer exosomes

The nano-flow cytometry method for the analysis of exosomes combines different fluorescent labelling of the vesicles. All experimental details can be found in S6 (Supp. Data).

#### CFSE and Violet staining of exosomes

Firstly, exosomes derived from SKBR3, MDA-MB-231 and MCF7 breast cancer cell lines were labelled with cell tracking reagents. From the same commercial source, a range of different colors are available (*i.e.,* referred to CellTrace^TM^ kits, Thermo Fisher Scientific), allowing to adjust the exosome staining for each experiment with combination of fluorophores. For exosome tracing, we used CellTrace™ CFSE (Ex 488 nm/Em 525 nm; Thermo Fisher Scientific, C34554) and CellTrace™ Violet (Ex 405 nm/Em 450 nm; Thermo Fisher Scientific, C34557). These cell-permeant succinimidyl ester dyes diffuse through the exosomal membrane and covalently label primary amines on proteins, providing a stable fluorescence signal for single-particle analysis.

In this study, CellTrace^TM^ CSFE proliferation kit (excitation 488 nm, emission 525 nm, ref. C34554, Thermo Fisher Scientific) and CellTrace^TM^ Violet proliferation kit (excitation 425 nm, emission 450 nm, ref. C34557, Thermo Fisher Scientific) were used for exosome staining (Fig. S5, panel A). Cell tracking reagents react covalently with proteins and due to their fluorescence emision they allow to separate the exosomes from background signals (Ender et al., 2020; Morales-Kastresana et al., 2017). The incubation parameters (exosome concentration, temperature, and time) for exosome labelling with cell tracking reagents, as well as the instrumental parameters of the nano-cytometer (channels gain, threshold, and range) were optimized. The optimal exosome concentration ranges from 10^8^ to 10^9^ particles · mL^−1^, according to NTA measurements. The CFSE/Violet concentration was optimized to 20 μmol · L^−1^, with 2 hours of incubation at 37°C. After incubation, all samples were diluted to 1 mL with HEPES buffer and stored at 4°C until measurement by nano-flow cytometry. Regarding the instrumental parameters, the gain of B525-FITC channel was set to 1,000 units, while V450-Violet to 500 units. Thresholds were adjusted manually at 700 units in both channels.

#### Analysis of membrane markers on CFSE-labelled exosomes

For the detection of surface membrane proteins on exosomes, the Violet vesicles (CellTrace Violet-labelled exosomes) were incubated with FITC-modified primary antibodies against tetraspanins CD9, CD63 and CD81 (Fig. S5, panel B). This assay allows to locate the exosomes (tetraspanin-containing vesicles) on the subpopulation of vesicles that were labelled with violet staining (protein-containing vesicles). The assay was done with 20 µL of exosomes (containing 8.60 x 10^7^ particles, according to NTA counting) derived from SKBR3 breast cancer cell lines directly labelled with 5 µL of FITC-modified antiCDX mouse monoclonal antibodies (being CDX either CD9, CD63 or CD81 biomarkers) for 45 min at RT. In parallel, non-labelled control samples (not incubated with the cell tracking reagent) and without antibodies were prepared with the same corresponding volumes of HEPES buffer. After incubation, all samples were diluted to 1 mL with HEPES buffer and stored at 4°C until measurement by nano-flow cytometry.

#### ALDH activity determination in breast cancer exosomes

To determine the ALDH activity on exosomes derived from SKBR3, MDA-MB-231 and MCF7 breast cancer cell lines, exosomes were labelled with the CellTrace CFSE (from now, named as CFSE-exosomes) and incubated with RP substrate (Fig. S5, panel C). This small, hydrophobic, and non-ionic molecule is able to cross the exosome lipidic bilayer membrane and react with the intra-exosome ALDH enzyme to produce resorufin (excitation 525 nm, emission 585 nm) by its esterase ability. The concentration of the substrate was optimized at 25 μmol L^-1^ with 2 hours at 25°C incubation. RP substrate stock was dissolved in DMSO, and ABD0305 inhibitor, in ethanol. Therefore, and to not adversely affect the biological activity of the exosomes, at least a 100-fold dilution in HEPES buffer from the stocks were used as working solutions. Besides ALDH, other esterase enzymes are also able to react with the substrate. Therefore, a specific ALDH inhibitor (ref. ABD0305) was used to indirectly determine the ALDH activity by inhibiting its contribution to the resorufin signal. The concentration of the inhibitor was optimized at 0.4 mmol L^-1^ with 2 hours at 37°C incubation, always prior to substrate incubation. The resorufin was detected with the Y610 detector, showing better results than Y585 detector. Additionally, to evaluate the effect of the storage temperature on the ALDH enzymatic activity, a comparative study was done with exosomes obtained from i) 5-days frozen (-21°C) or ii) freshly obtained (<12h) and non-frozen (4°C) cell culture supernatants.

### 3.11 Statistical analysis

The statistical analyses were performed using GraphPad Prism v10 (CA, US). All experiments were done at least by duplicate. In the flow cytometry experiments, at least 10,000 cells or vesicles were recorded per sample.

### 3.12 Safety considerations

All works were performed in a Biosafety cabinet, and all material was decontaminated by autoclaving or disinfected before discarding following U.S. Department of Health and Human Services guidelines for level 2 laboratory Biosafety (CDC, 2020).

## 4. Results and discussion

### 4.1 Expression levels of ALDH isoforms in breast cancer cell lines

To establish ALDH expression profile across breast cancer molecular subtypes, we analyzed three representative cell lines: SKBR3 (HER2-enriched), MDA-MB-231 (triple-negative), and MCF7 (luminal A), classified according to the PAM50 gene expression assay.(Prat et al., 2012). We assessed class 1A isoforms (ALDH1A1, ALDH1A2, ALDH1A3), ALDH1B1 (class 1B), ALDH2 (class 1) and ALDH3A1 (class 3) by capillary electrophoresis immunoassay (Figure 2).

**Figure 2.**
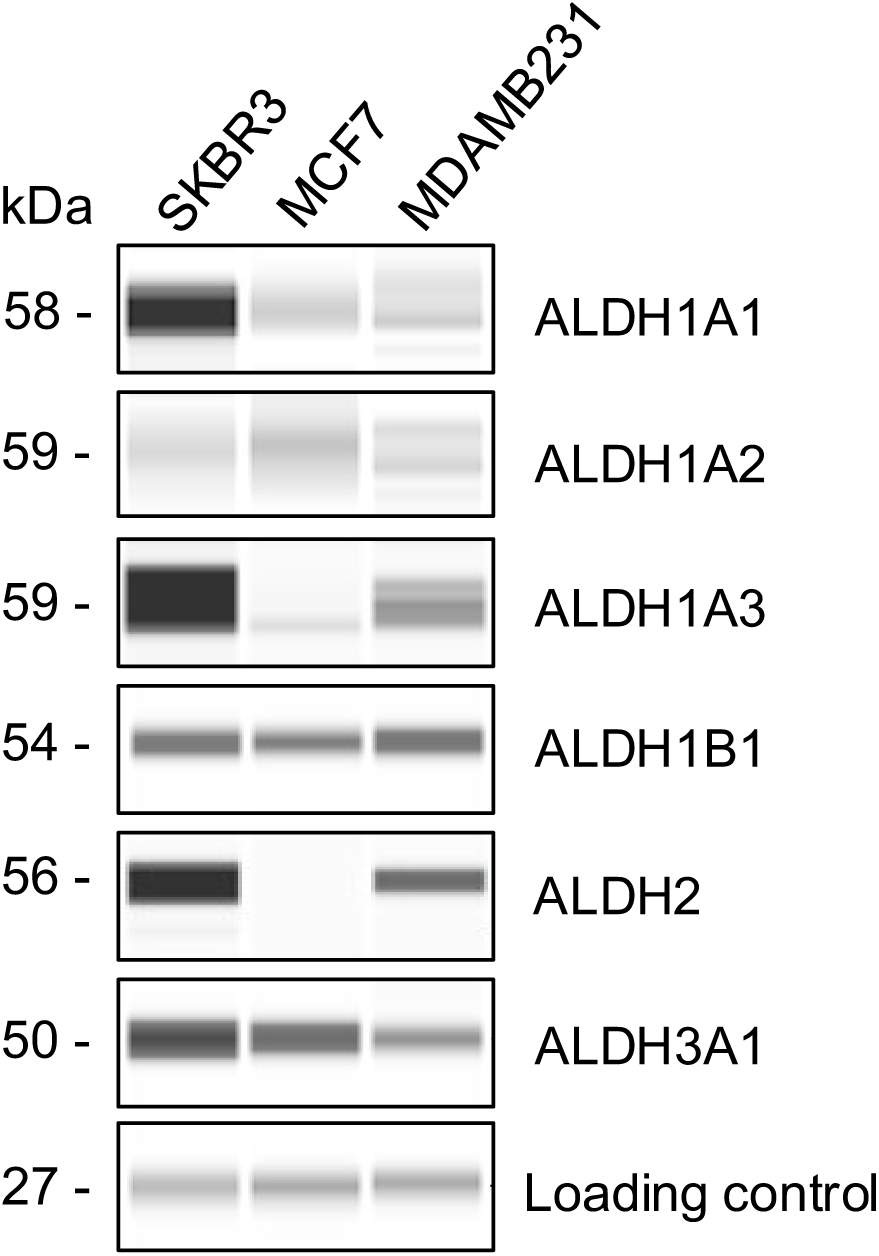
Capillary electrophoresis immunoassay showing the endogenous amount of ALDH isoforms in SKBR3, MDA-MB-231 and MCF7 breast cancer cell lines. Comparisons can be made across all cell lines for a single isoform, but direct comparisons between different isoforms within the same cell line are not possible due to antibody affinity differences and consequent variations in experimental conditions. The image was cropped to show the signals corresponding to ALDHs only.

All three cell lines expressed ALDH1B1, while other isoforms varied in abundance. Among class 1A isoforms, ALDH1A3 was highest in SKBR3 and MDA-MB-231 and barely detectable in MCF7. ALDH1A1 was also elevated in SKBR3 relative to the other lines, whereas ALDH1A2 was low across all models. For class 3, ALDH3A1 was consistently expressed without clear subtype specificity. Collectively, HER2-enriched and triple-negative breast cancer cells show stronger ALDH1A3 (and, in SKBR3, ALDH1A1) expression compared with luminal A cells, consistent with reports linking these isoforms to aggressive, therapy-resistant phenotypes (Pequerul et al., 2025).

### 4.2 Intracellular staining of ALDH in breast cancer cells

To validate these findings at the single-cell level, we analyzed intracellular ALDH1A3 by immunofluorescence and flow cytometry (Figure 4). Cell permeabilization and integrity were confirmed using anti-vinculin staining, which yielded >90% positive cells across all lines (99.6% in SKBR3, 99.0% in MDA-MB-231, and 92.5% in MCF7; Supplementary Data S4, Figure S2). Secondary antibody specificity was verified in samples stained without primary antibody, showing <2% positivity in SKBR3 and MDA-MB-231 and 6% in MCF7.

Consistent with the WES data, the vast majority of SKBR3 and MDA-MB-231 cells were ALDH1A3-positive (98.8% and 89.5%, respectively), whereas MCF7 showed a lower fraction of ALDH1A3-positive cells (≈82%), indicating that up to one-fifth of the population lacked detectable expression. These results align with recent studies showing that ALDH1A3 is broadly expressed within HER2-enriched and triple-negative cells, while its expression is more heterogeneous in luminal A cells, supporting a subtype-specific pattern of ALDH1A3 regulation (Pequerul et al., 2025).

### 4.3 Fluorometric determination of ALDH enzymatic activity in breast cancer cells

Next, we quantified ALDH enzymatic activity in whole-cell lysates from SKBR3, MDA-MB-231, and MCF7 using a fluorometric assay that monitors NADH production during aldehyde oxidation. Hexanal was used as substrate, and assay conditions (buffer/ionic strength and cofactor concentrations) were optimized to favor ALDH1A3 detection relative to other 1A isoforms, based on reported kinetic preferences (Jiménez et al., 2024; Pequerul et al., 2025, 2022, 2020).

Consistent with the ALDH1A3 protein levels (Figure 3) and intracellular staining (Figure 4), all three lines exhibited detectable activity in fresh lysates, ranging 0.29–0.95 mU·mg⁻¹ protein (Table 1). As expected for labile dehydrogenase activity, a single freeze–thaw cycle produced a marked reduction in signal. The magnitude of activity was comparable to values reported for ALDH1A3-expressing A172 glioblastoma cells (∼1.79 mU·mg⁻¹) (Jiménez et al., 2024), supporting the assay’s physiological relevance.

**Figure 3.**
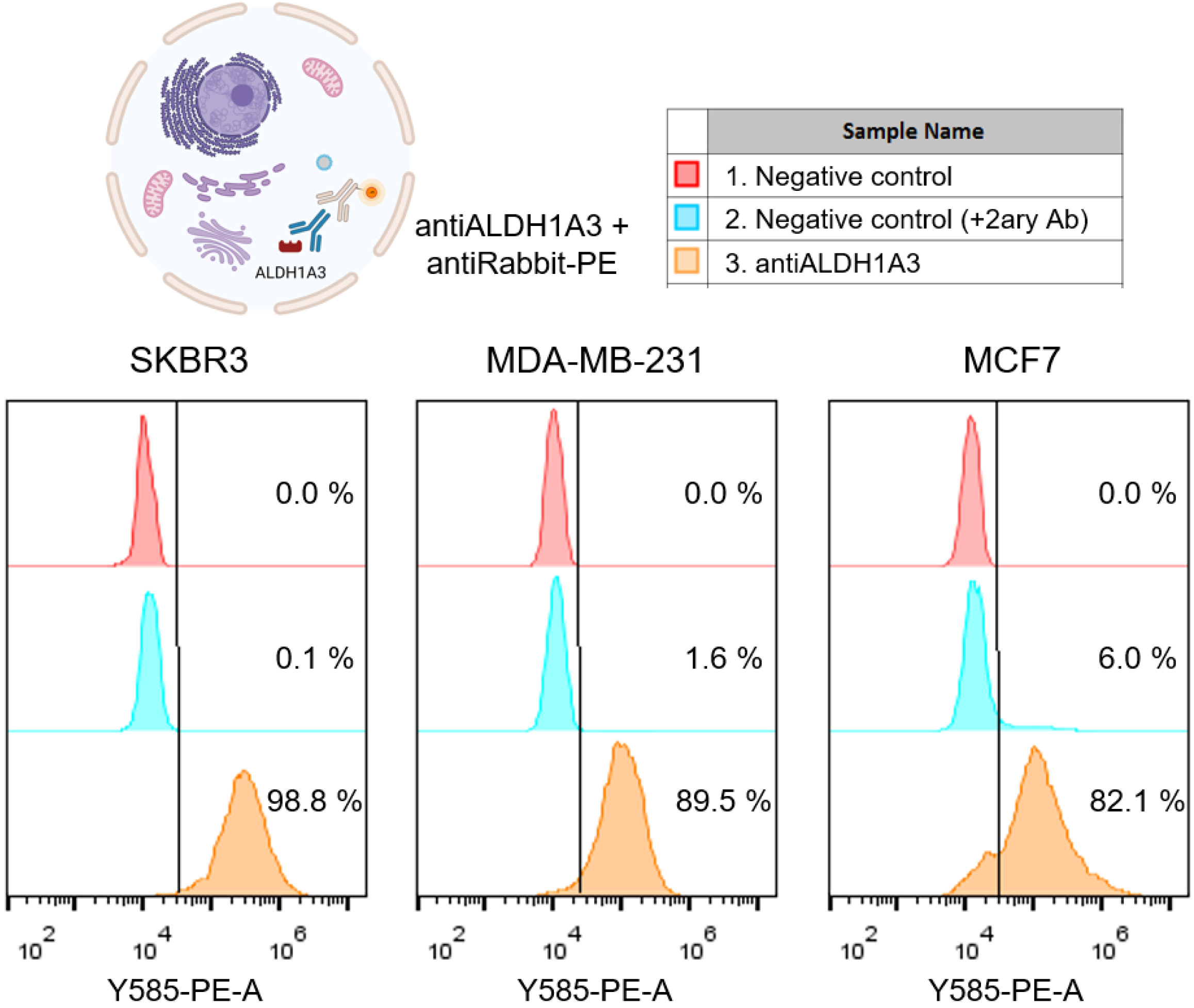
Histograms from intracellular staining of cells from SKBR3, MDA-MB-231 and MCF7 breast cancer cell lines with anti-ALDH1A3 antibodies, labelled with PE-modified anti-rabbit secondary antibodies. In red color, negative controls of the experiment, only cells; in blue color, control with cells only incubated with secondary antibodies; in orange color, cells with antiALDH1A3 antibodies plus labelled secondary antibodies.

**Table 1.** ALDH activity in (mU·mg⁻¹ protein) in whole-cell lysates of breast cancer cell lines (SKBR3, MDA-MB-213 and MCF7), measured by a NADH-based fluorometric assay using hexanal as substrate under ALDH1A3-preferential assay conditions^1^.

| SAMPLE | SPECIFIC ALDH<br>ACTIVITY<br>($\text{mU} \cdot \text{mg}^{-1}$) |
| --- | --- |
| SKBR3<br>(FRESH) | $0.95 \pm 0.04$ |
| SKBR3<br>(FROZEN) | $0.61 \pm 0.04$ |
| MDA-MB-231<br>(FROZEN) | $0.29 \pm 0.04$ |
| MCF7<br>(FROZEN) | $0.84 \pm 0.12$ |
| SKBR3 EXOSOMES<br>(FROZEN) | $0.009 \pm 0.002$ |

<sup>1</sup>Fluorescence of the NADH reaction product was monitored at 460 nm with excitation at 340 nm in 50 mM HEPES, 50 mM $\text{MgCl}_2$ , 5 mM DTT, pH 8.0 and 25°C, using 250 $\mu\text{M}$ hexanal as substrate. Data is presented as the mean ± standard deviation from triplicate measurements.

Together, these data confirm functional ALDH activity consistent with ALDH1A3 abundance and underscore the importance of fresh sample handling for reliable measurement.

In parallel, the ALDH activity was also determined in exosomes derived the three cell lines. Nevertheless, only exosomes from SKBR3 cell line could be quantified (0.009 mU/mg protein), while MDA-MB-231 and MCF7 remain under the LOD of the method.

### 4.4 Nano-Flow cytometry studies of breast cancer exosomes

Having established ALDH1A3 expression and enzymatic activity in breast cancer cells, we then examined whether this isoform could also be detected in exosomes released by these cells. As expected, ALDH1A3 was not detected in the exosomal membrane by ELISA (Supplementary Figure S4), consistent with its cytosolic localization. However, ALDH activity was clearly observed within the exosomal cargo of SKBR3-derived vesicles, providing functional evidence that the enzyme is encapsulated and remains catalytically active after secretion.

Previously, our group demonstrated the use of intrinsic enzymatic activity as a biomarker for exosome detection, specifically, alkaline phosphatase activity in exosomes purified from the human fetal osteoblast (hFOB 1.19) cell line and from human serum of healthy individuals (Moura et al., 2022a; Sappia et al., 2019). Based on this strategy, we explored ALDH1A3 activity as a functional biomarker of ALDH1A3-positive exosomes from breast cancer cells.

Exosomes were analysed in suspension by nano-flow cytometry, a method that enables single-vesicle detection without the need for immobilization on microparticles or capture beads. This is a major methodological advantage, as it allows exosomes to be examined directly from purified preparations, preserving their native size distribution and reducing handling artifacts. (Ender et al., 2020; Hoen et al., 2012).

For discrimination from background noise, exosomes were fluorescently labeled using cell-permeant amine-reactive tracers CellTrace CFSE and CellTrace Violet, which covalently bind to intravesicular and membrane-associated proteins. These dyes provided bright and stable signals suitable for single-vesicle detection. SKBR3-, MDA-MB-231–, and MCF7-derived exosomes were successfully detected using this approach, hereafter referred to as CFSE-exosomes and Violet-exosomes, respectively (Figure 4A).

**Figure 4.**
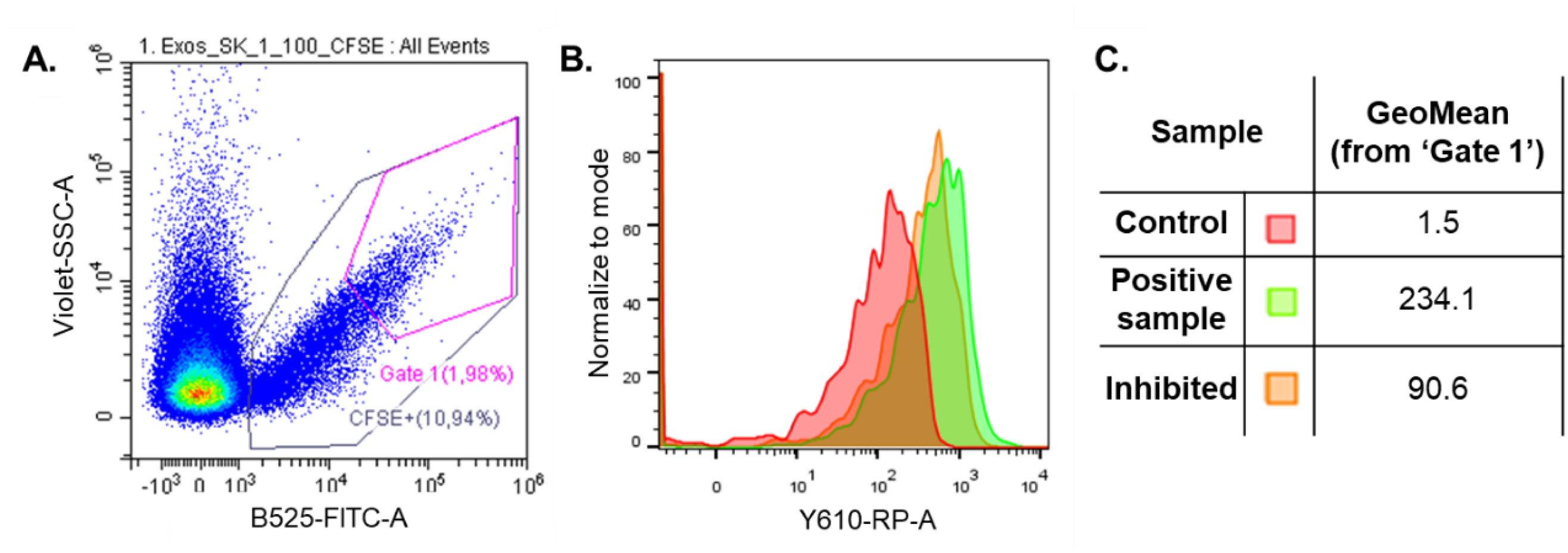
Analysis of the resorufin propionate reaction for ALDH activity detection on exosomes by nano-flow cytometry. On the Panel A, dot plot representation of CFSE-labelled SKBR3 exosomes as control sample, with total CFSE positive and Gate 1 subpopulations selected. On Panel B, histogram representation of the resorufin signals from the three SKBR3 samples. On Panel C, geometric mean of the Y610-RP signals from ‘Gate 1’.

To assess ALDH activity, CFSE-labeled exosomes were incubated with the resorufin propionate (RP) substrate, whose reaction with active ALDH produces a distinct fluorescence signal. The dual-fluorescence design (CFSE + RP) constitutes a flexible analytical platform, enabling simultaneous detection of total vesicle counts and enzymatic activity within the same population. The spectral separation between CFSE (emission ∼517 nm) and RP (emission ∼585 nm) allows concurrent acquisition in FITC and PE channels without significant overlap, facilitating robust multiparametric analysis by standard flow cytometry.

#### Experimental design of ALDH determination of tracer-labelled exosomes by nano-flow cytometry

To ensure the reliability and sensitivity of ALDH activity detection in exosomes, we first optimized the fluorescent tracer labelling protocol for nano–flow cytometry analysis. Several experimental variables were systematically evaluated (including exosome concentration, dye concentration, incubation time and temperature, and instrument gain and threshold settings) as detailed in Supplementary Data S7 and Figures S5–S9.

Under the optimized conditions, CellTrace CFSE and CellTrace Violet provided bright and stable labeling suitable for single-vesicle detection. Optimal staining was achieved with 20 μmol·L^−1^ dye and 2 h incubation at 37°C with gentle shaking, using an exosome suspension of approximately 10^7^ particles·mL^−1^ as determined by NTA (Fig. S7). These optimized parameters enabled reproducible detection of exosomes as discrete fluorescent events, minimizing background noise and signal variability. Fluorescent tracer–labeled exosomes were subsequently used to identify tetraspanin enriched vesicle populations, defined as *Gate 1* subpopulation (Fig. S11). These results confirm that the optimized nano–flow cytometry setup reliably detects and discriminates exosome subpopulations in solution, providing a robust analytical basis for the subsequent detection of enzymatic activity.

To establish conditions for ALDH activity quantification, we next adapted a fluorogenic substrate system to exosomal analysis. The standard ALDEFLUOR™ assay, based on BODIPY–aminoacetaldehyde and DEAB, lacks isoform specificity and spectral compatibility with CFSE. Therefore, we selected resorufin propionate (RP), a small non-ionic substrate recently characterized as an ALDH1A3-compatible fluorogenic probe (Ceylan et al., 2022; unpublished results). RP can diffuse across lipid membranes and be cleaved by ALDH through its esterase activity, yielding the fluorescent product resorufin (λ_exc = 560 nm; λ_em = 583 nm) with a high quantum yield (Φ = 0.75). To ensure reaction specificity, the selective ALDH1A3 inhibitor ABD0305 (Pequerul et al., 2025) was used as a control to discriminate ALDH-dependent from non-specific esterase signals. Key parameters, including substrate and inhibitor concentration, temperature, and incubation time, were optimized as described in Supplementary Data S7.2. The final assay conditions were 25 μmol·L⁻¹ RP for 2 h at 25 °C, and 0.4 mmol·L⁻¹ ABD0305 for 2 h at 37 °C (Figure S11). The fluorescent product resorufin was detected in the Y610-RP channel of the flow cytometer.

Exosome samples, including positive controls (RP-treated), negative controls (unstained), and inhibitor-treated preparations, were analyzed within the Gate 1 population, corresponding to tetraspanin-enriched vesicles (Figure S10). Fluorescence was recorded in the Y610-RP channel, and the geometric mean intensity of resorufin emission was used as a quantitative measure of ALDH activity. As illustrated for SKBR3-derived exosomes (Figure 5), the RP-treated sample displayed a clear fluorescence shift compared with the negative control and the ABD0305-treated sample, which was pre-incubated with the selective ALDH1A3 inhibitor before substrate addition. The geometric mean fluorescence of the positive sample (234.1) was approximately 2.6-fold higher than that of the inhibited sample (90.6), confirming that a substantial portion of the RP signal originated from ALDH catalysis within the exosomal lumen. By normalizing the geometric mean of the positive control to 100% total activity, ABD0305 inhibition reduced the signal to 38.7%, indicating that ALDH1A3 accounts for approximately 61% of the total esterase-like activity detected in SKBR3 exosomes. Together, these results provide direct functional evidence that enzymatically active ALDH1A3 is present as cargo within breast cancer–derived exosomes and demonstrate the feasibility of quantifying isoform-specific enzymatic activity by nano–flow cytometry.

**Figure 5.**
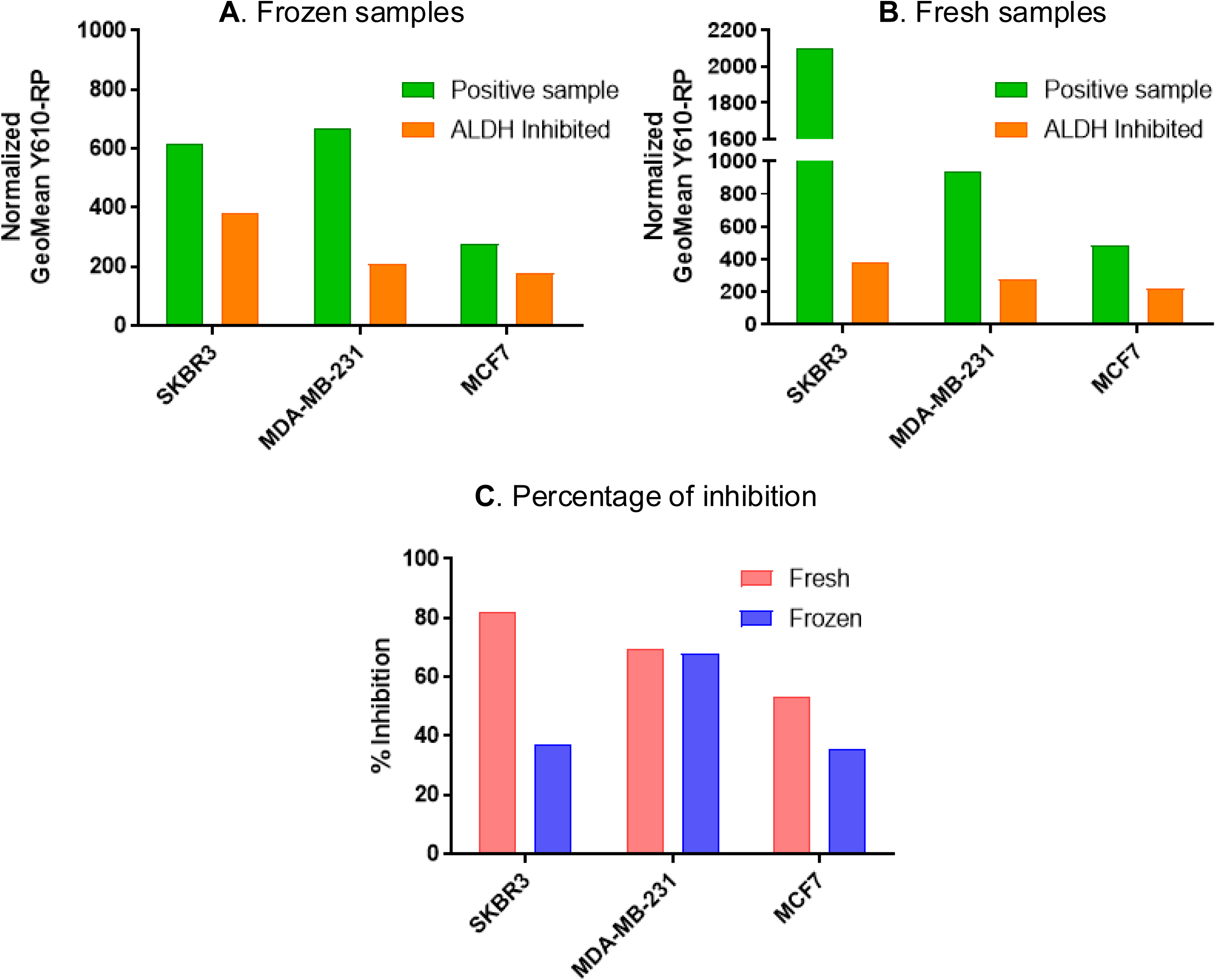
Effect of the storage temperature on the ALDH activity of the exosomes. A, bar graph representation geometric mean on Y610-RP-A channel (normalized with negative controls, respectively) of exosomes fresh samples from SKBR3, MDA-MB-231 and MCF7 breast cancer cell lines. B, bar graph from frozen samples from the three cell lines. Blue bars: control samples; green bars: positive samples; orange bars: inhibited samples. C, the percentage of inhibition is calculated for frozen and fresh samples by the normalized difference of positive versus inhibited samples.

#### Determination of ALDH1A3 activity in breast cancer-derived exosomes Stability of exosomal ALDH activity and impact of sample freezing

To evaluate the stability of ALDH enzymatic activity in breast cancer–derived exosomes and assess the influence of storage conditions, we analyzed exosome preparations from SKBR3, MDA-MB-231, and MCF7 cells using the same nano–flow cytometry protocol described above. In agreement with fluorometric assays, SKBR3-derived exosomes exhibited the highest ALDH activity, whereas MDA-MB-231 and MCF7 exosomes showed lower but detectable activity levels. Unexpectedly, when a new batch of SKBR3 exosomes was analyzed, the resorufin fluorescence signal was approximately threefold higher than that observed in the previous experiment (Fig 5A). Because the standard protocol for exosome isolation involves freezing of the supernatants and the purified vesicles at −2 °C (Moura et al., 2022b, 2020), we hypothesized that freeze–thaw cycles might compromise ALDH activity.

To test this, exosomes were freshly isolated from culture supernatants within 4 h following the procedure in Supplementary Data S2, and ALDH activity was measured under the same experimental conditions. As shown in Figure 5, freeze–thawing markedly reduced ALDH activity across all samples, and this reduction was most pronounced for SKBR3-derived exosomes. Moreover, the inhibitory effect of ABD0305 was stronger in fresh samples than in frozen ones, further indicating that enzyme integrity and inhibitor accessibility are affected by storage. Quantitative analysis (Figure 5C) revealed that the ALDH1A3 contribution, estimated from ABD0305-mediated inhibition, decreased from 81.8 % to 37.1 % in SKBR3 exosomes after freezing, and from 53.0 % to 35.6 % in MCF7 exosomes, while remaining relatively stable in MDA-MB-231 exosomes (69.6 % to 67.9 %). Repetition of the SKBR3 experiments with independent fresh preparations confirmed the adverse effect of freezing on exosomal ALDH activity. These findings underscore that exosomal enzymatic activity is highly sensitive to storage conditions, highlighting the need for fresh sample processing when assessing functional biomarkers such as ALDH1A3 activity.

## 5. Conclusions

This study establishes a novel nano–flow cytometry–based method to detect and quantify ALDH1A3 enzymatic activity within breast cancer–derived exosomes. Exosomes represent an attractive source of biomarkers for liquid biopsy applications, offering minimally invasive access to molecular information otherwise obtainable only through tumor tissue. However, exploring the functional proteome within exosomal cargo remains challenging because the vesicle membrane must be preserved to maintain enzymatic integrity, while still allowing small substrates to diffuse inside. Our approach successfully overcame this limitation by enabling the detection of intravesicular ALDH1A3 activity without the need for membrane disruption. To our knowledge, this is the first demonstration of intra-exosomal enzymatic activity of ALDH1A3.

The key features of this assay include: (i) the use of cell-permeant fluorescent tracers for stable exosome labeling compatible with antibody staining, and (ii) the application of fluorogenic substrates capable of crossing the exosomal membrane to report enzyme activity directly through their catalytic conversion. Using RP as substrate and the ALDH1A3-selective inhibitor ABD0305, we were able to confirm the presence and functionality of this isoform in exosomal cargo and to quantify its relative contribution to total esterase-like activity.

While promising, the substrate specificity of RP remains limited, and future efforts should focus on developing new fluorogenic probes optimized for ALDH1A3 and compatible with nano–flow cytometry multiplexing. The spectral overlap encountered in triple-labeling assays also indicates that fluorophore combinations and compensation strategies will need refinement to expand the analytical versatility of this platform. Importantly, our data show that freeze–thaw cycles markedly reduce ALDH1A3 activity in exosomal samples, emphasizing that enzyme stability and storage conditions must be carefully controlled for reliable biomarker quantification. Consequently, this limitation underscores a key challenge in developing sensing strategies that rely on intrinsic ALDH1A3 activity in exosomes.

In summary, this work provides a proof of concept for detecting functional enzymatic biomarkers in intact exosomes and underscores the potential of nano–flow cytometry combined with fluorogenic substrates as a sensitive analytical approach for extracellular vesicle research and liquid biopsy applications.

## 6. Authors contributions

**A. Pallarès-Rusiñol:** Investigation, Validation, Writing – original draft. **R. Pequerul**: Investigation, Validation, Writing – original draft. **L. Costa-Sastre**: Investigation. **M. Tuxans**: Investigation. **A. Constantinescu**: Methodology, Validation. **M. Perez-Alea**: Methodology, Validation, Writing – revision. **M. I. Pividori**: Supervision, Funding acquisition. **J. Farrés**: Conceptualization, Methodology, Supervision, Writing – revision, Funding acquisition. **M. Martí:** Conceptualization, Methodology, Supervision, Writing – revision.

## 7. Acknowledgements

This work was partially funded by the Spanish Ministerio de Ciencia e Innovación (Agencia Estatal de Investigación, grant number PID2019-106625RB-I00), Ministerio de Ciencia, Innovación y Universidades (PID2023-150696NB-I00 /MCIU/ AEI / 10.13039/501100011033 /FEDER, UE). A. Pallares-Rusiñol acknowledges the Ministry of Universities (Grant FPU16/01579). R. Pequerul received financial support from Advanced BioDesign through a research contract agreement with the Universitat Autònoma de Barcelona. M. ernuz is acknowledged for providing exosome samples. Also, ICTS “NAN IOSIS” NTA analysis service of *Institut de Ciència dels Materials de Barcelona* and Service of Microscopy of *Universitat Autònoma de Barcelona* are gratefully acknowledged.

---

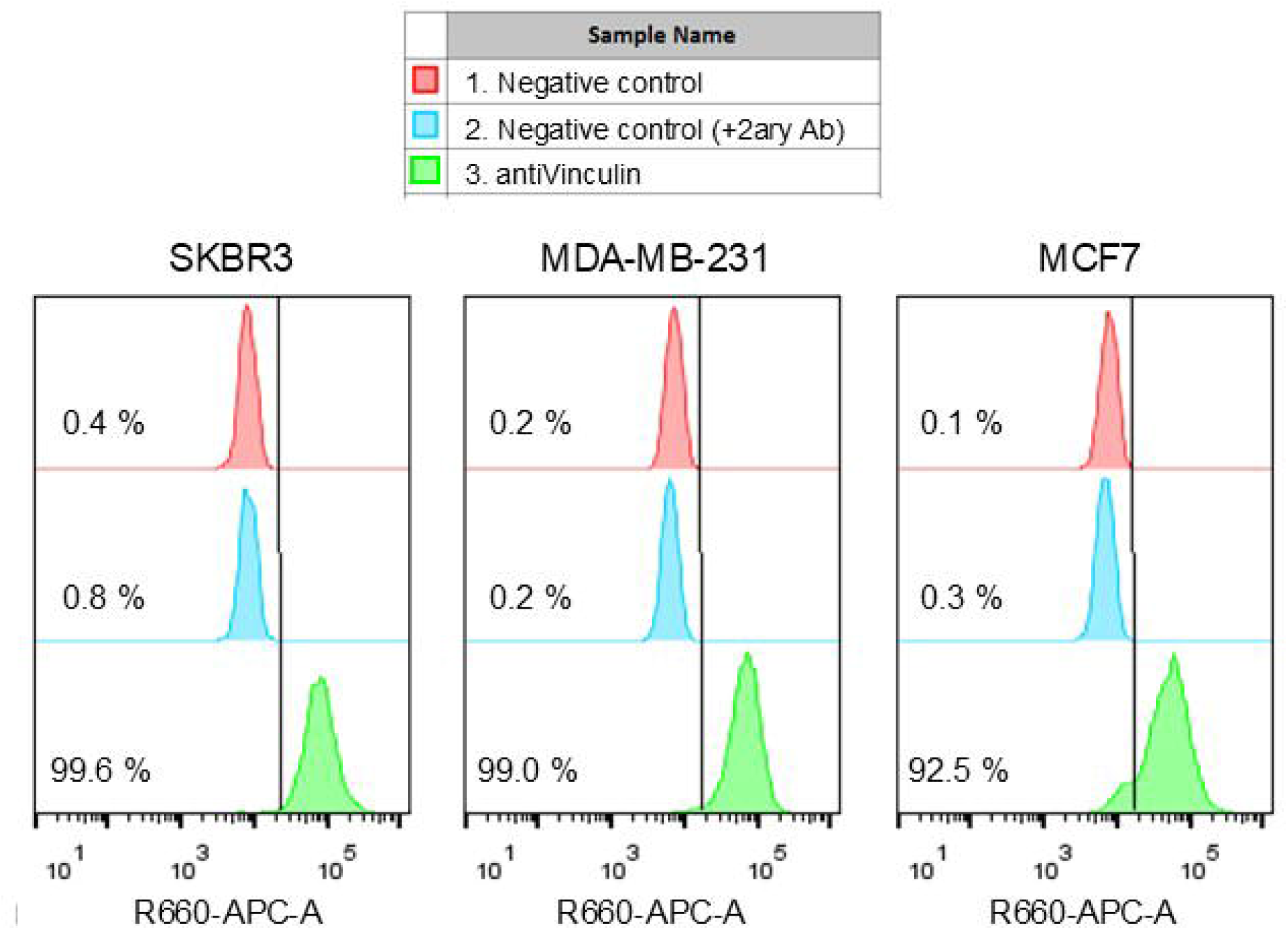

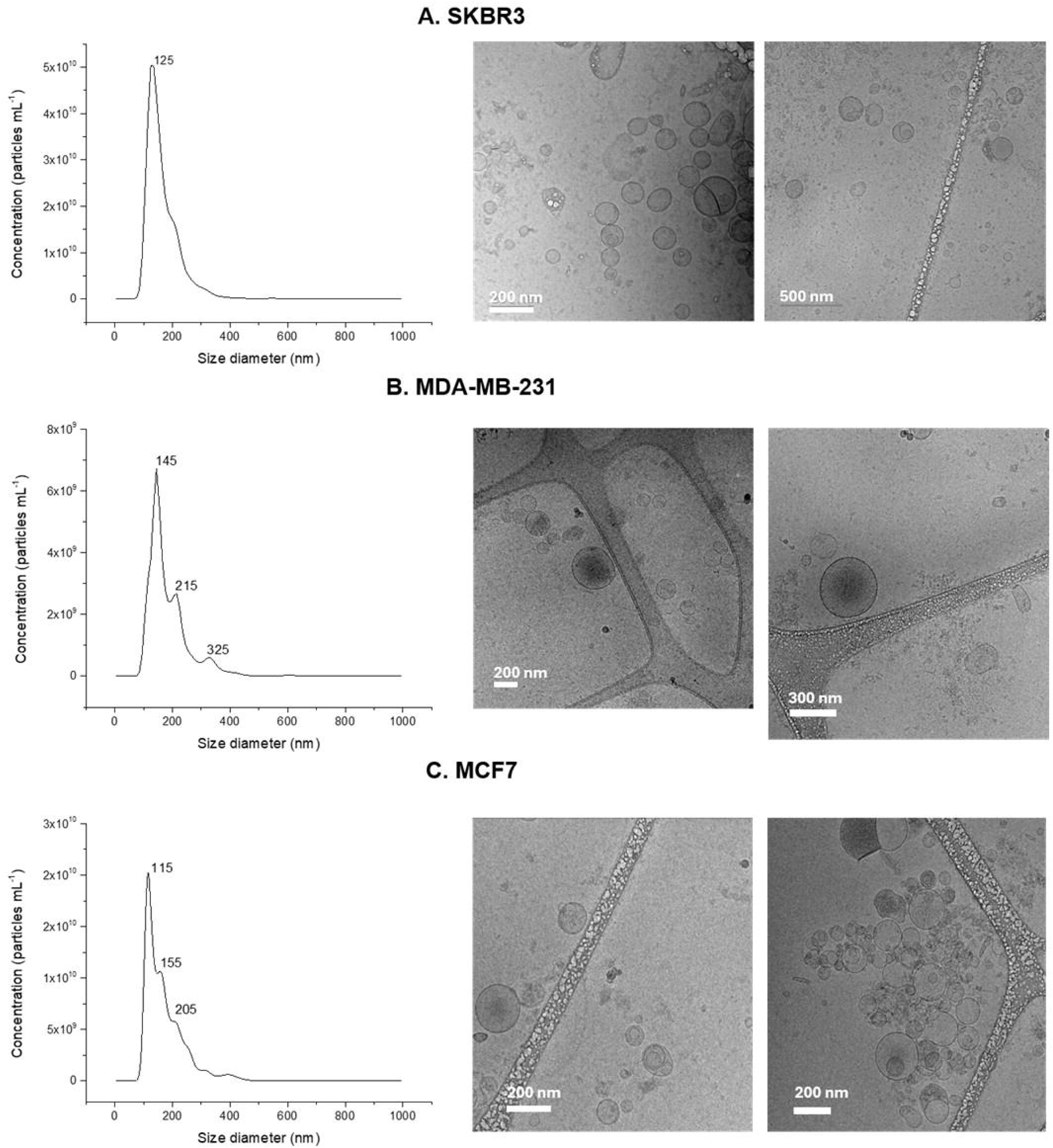

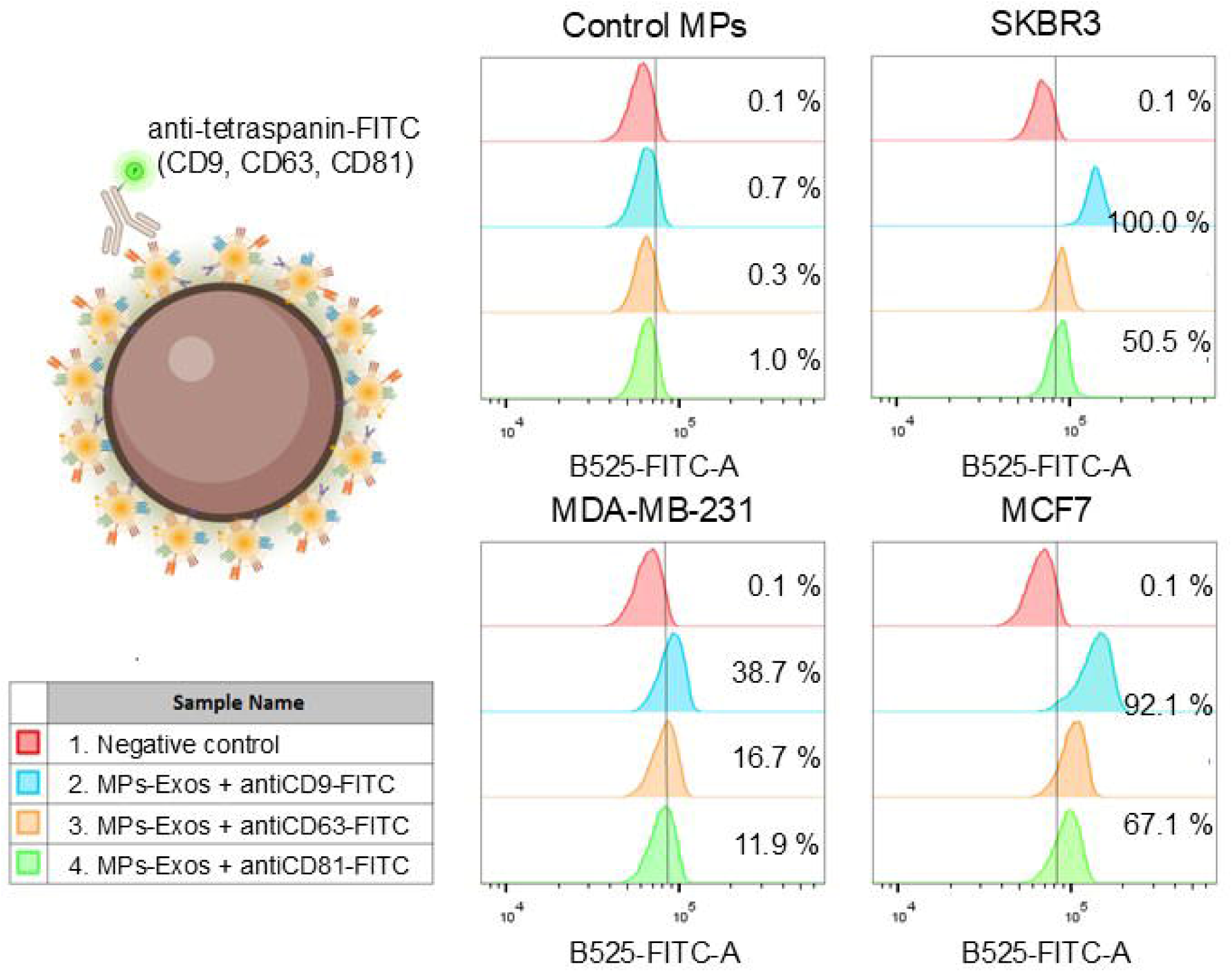

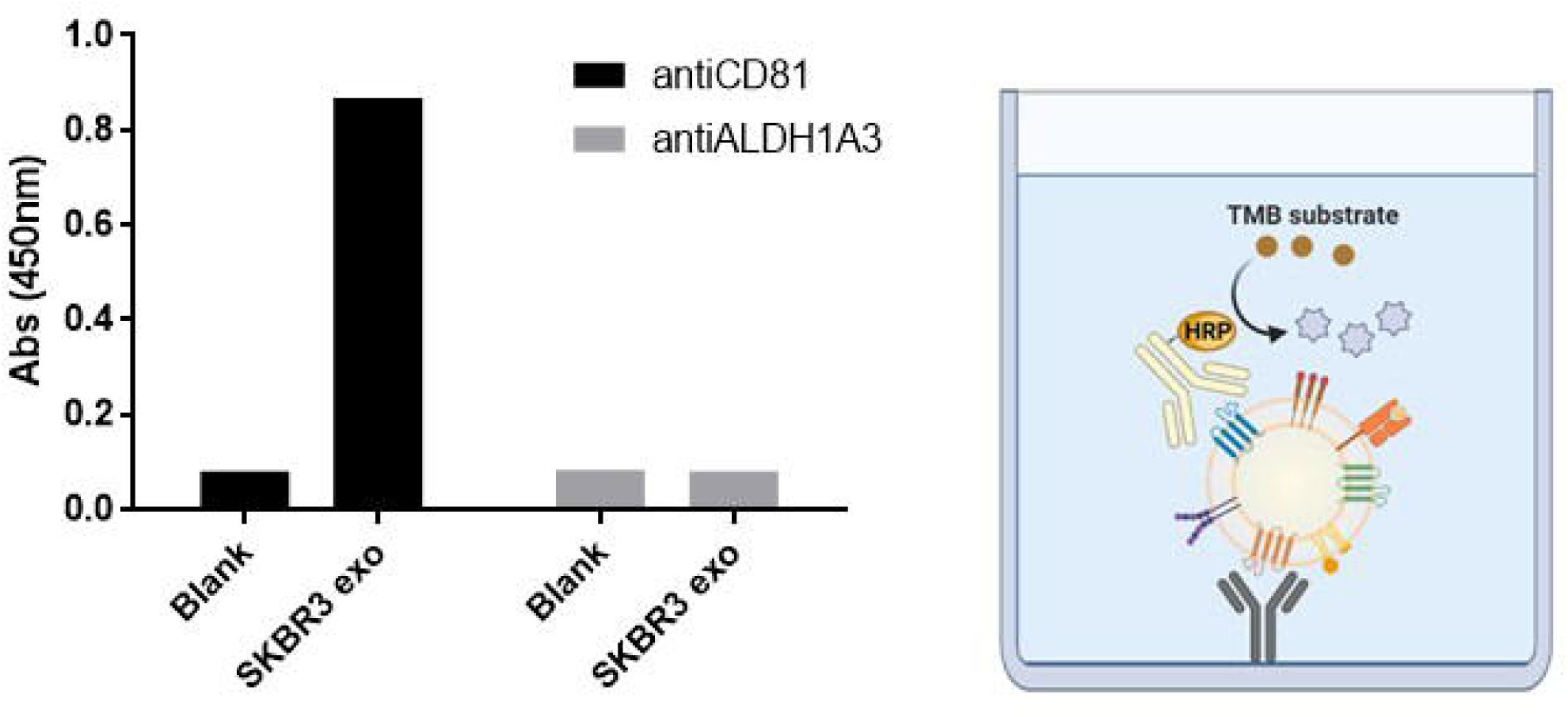

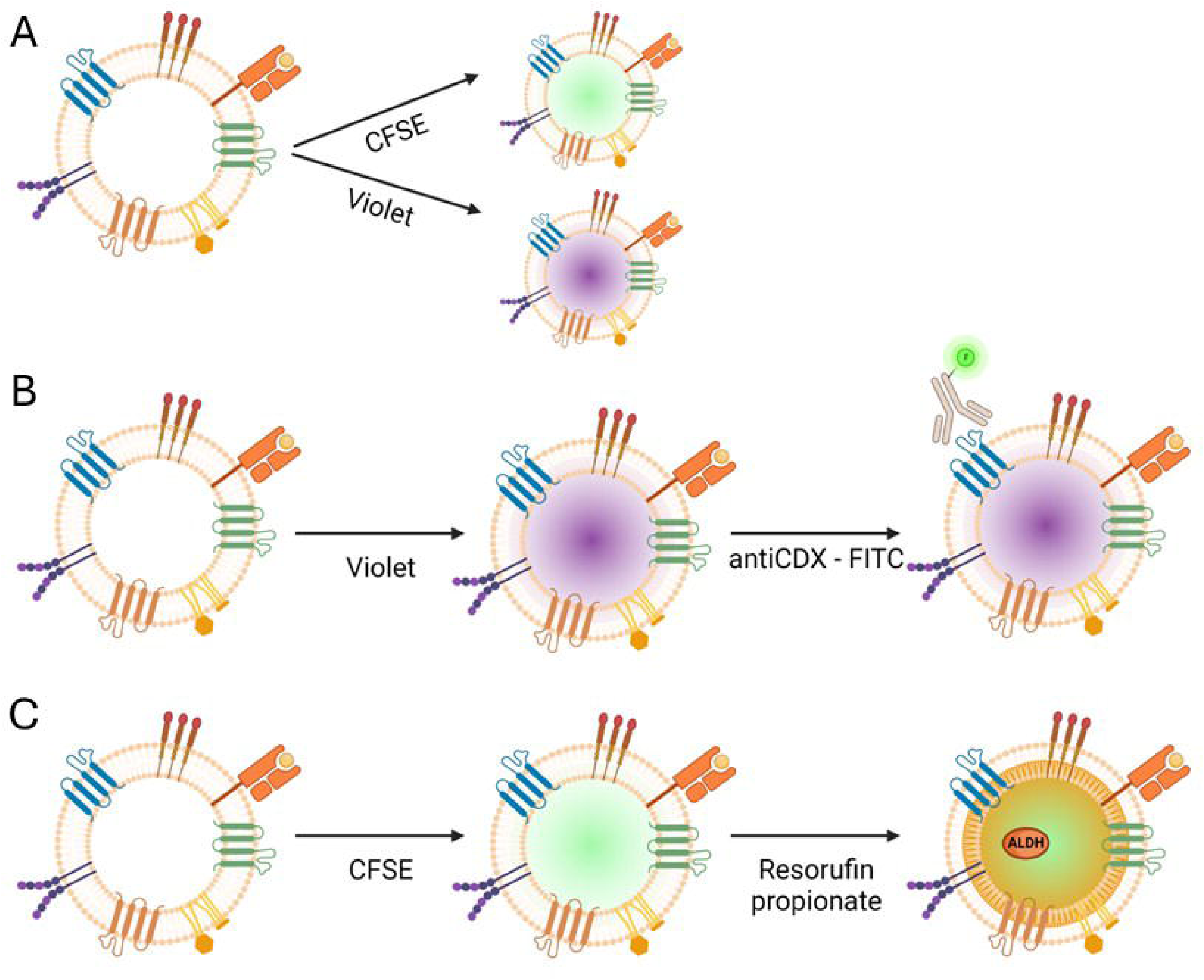

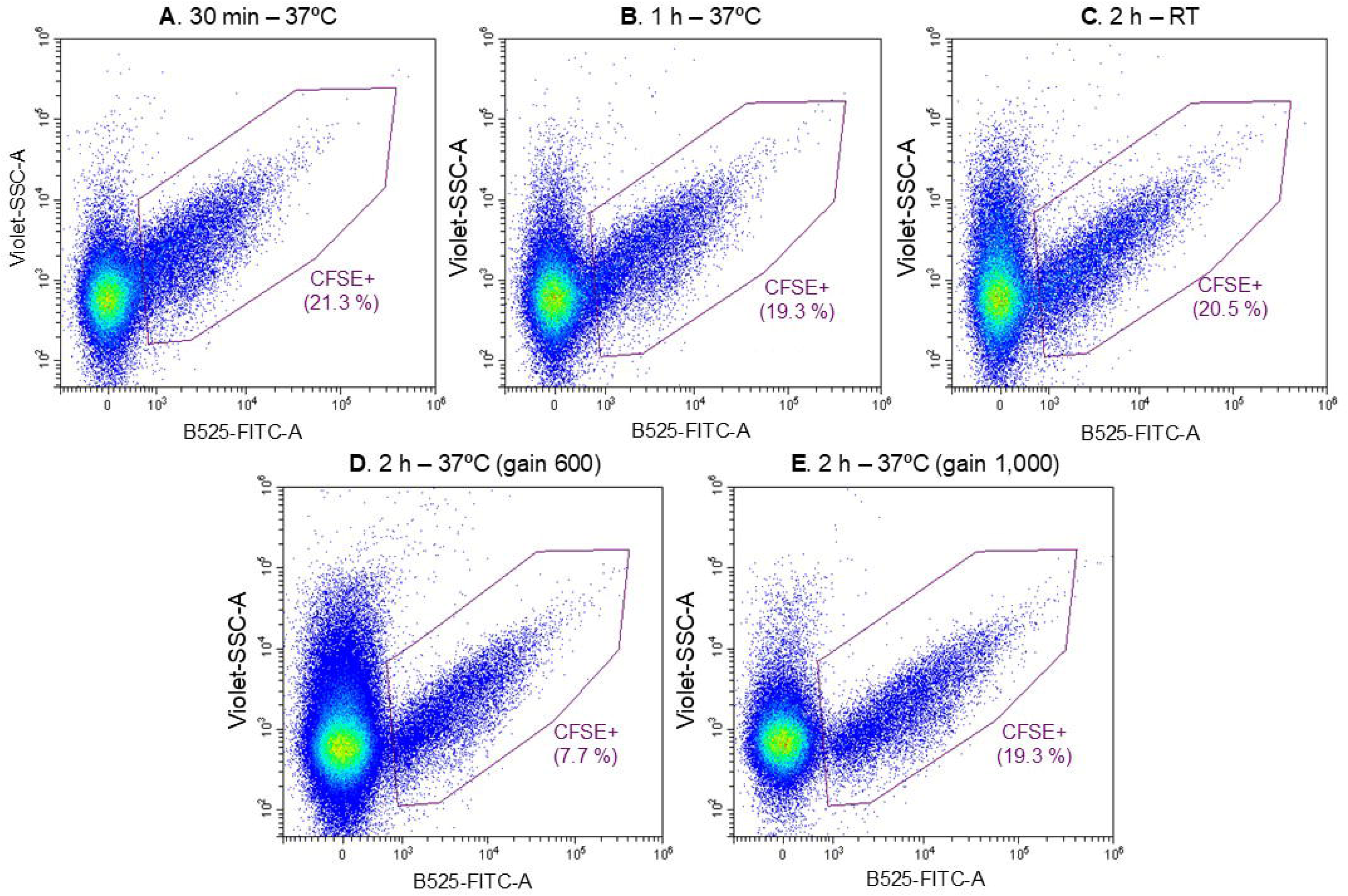

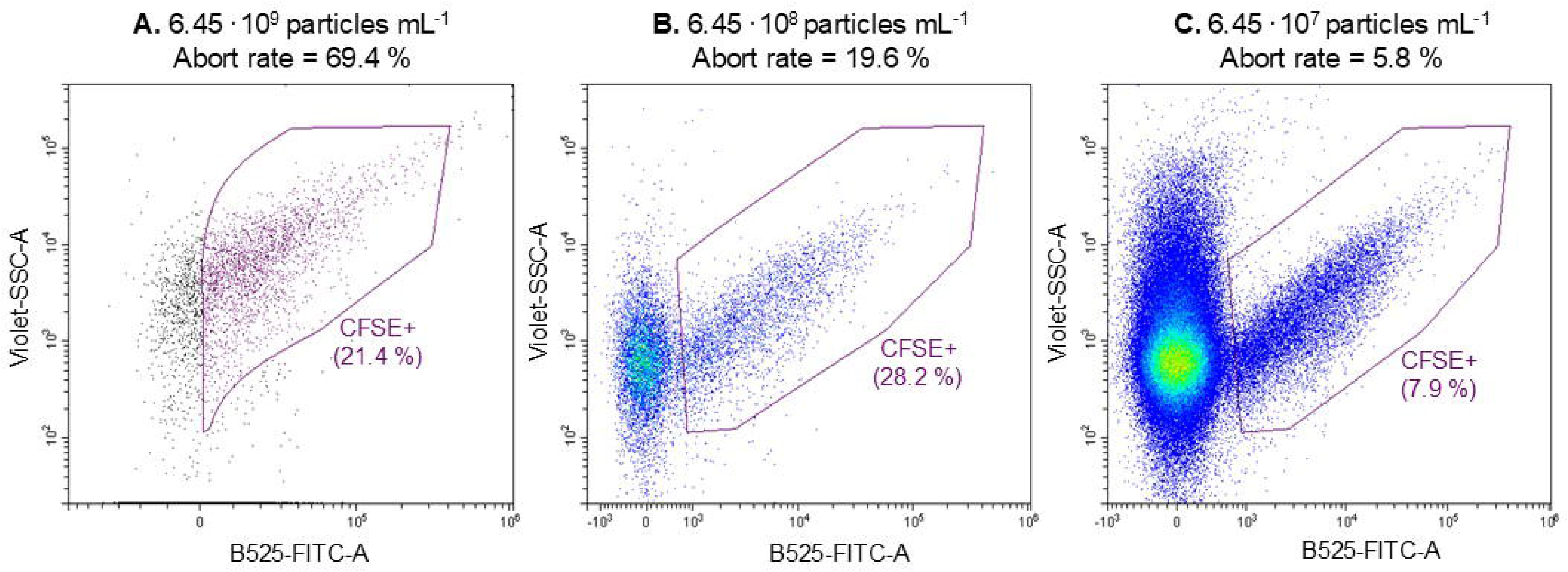

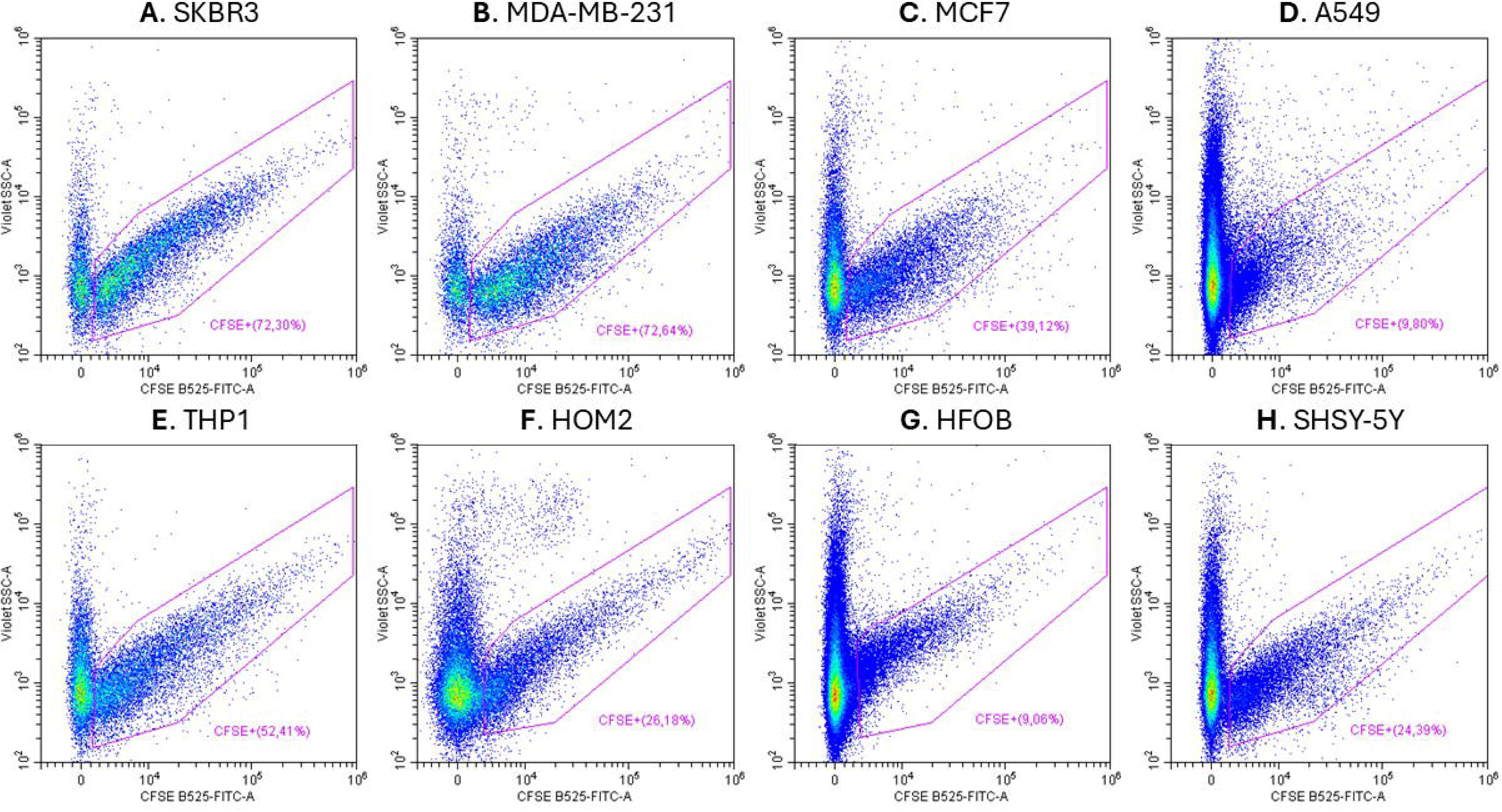

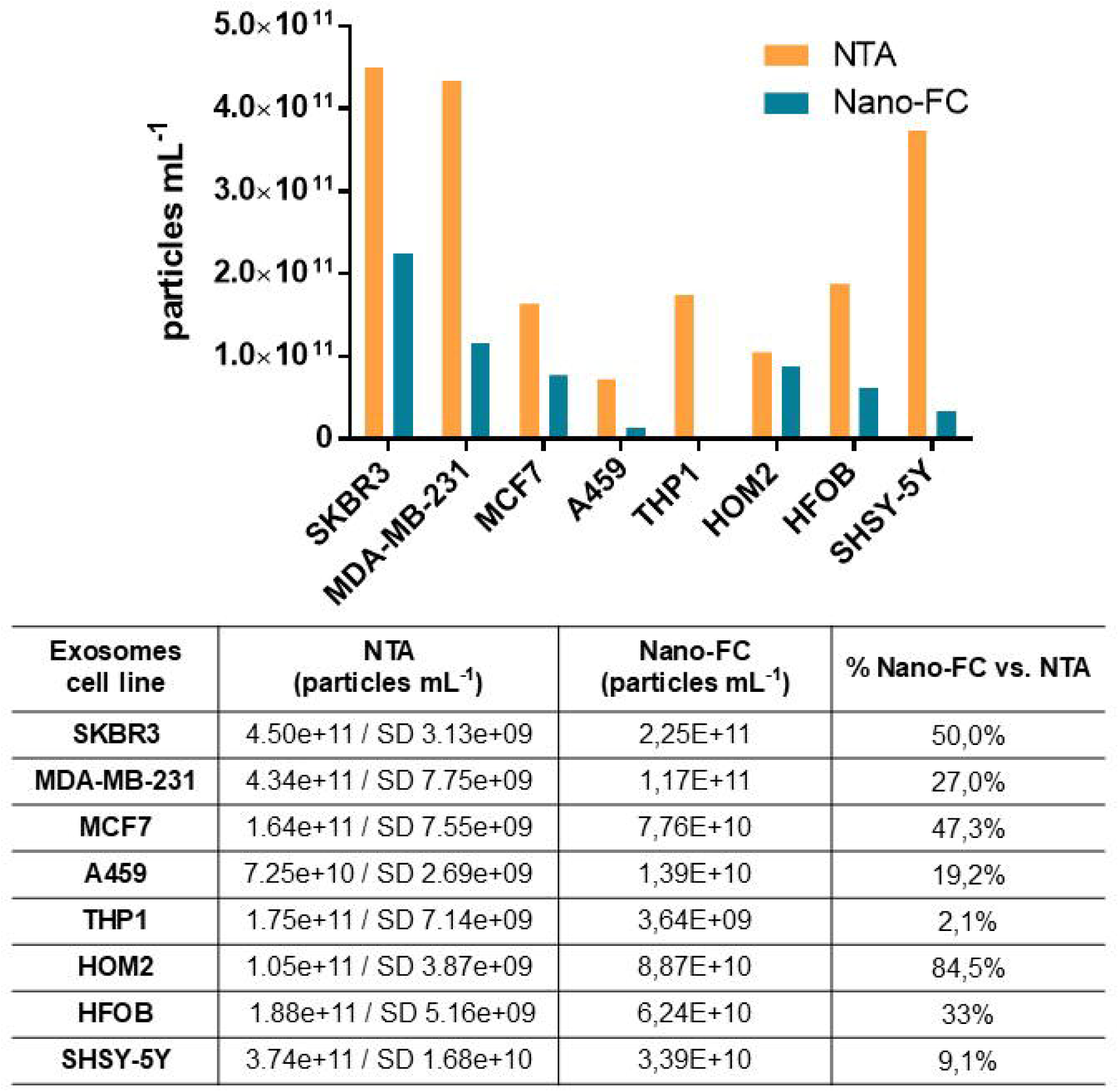

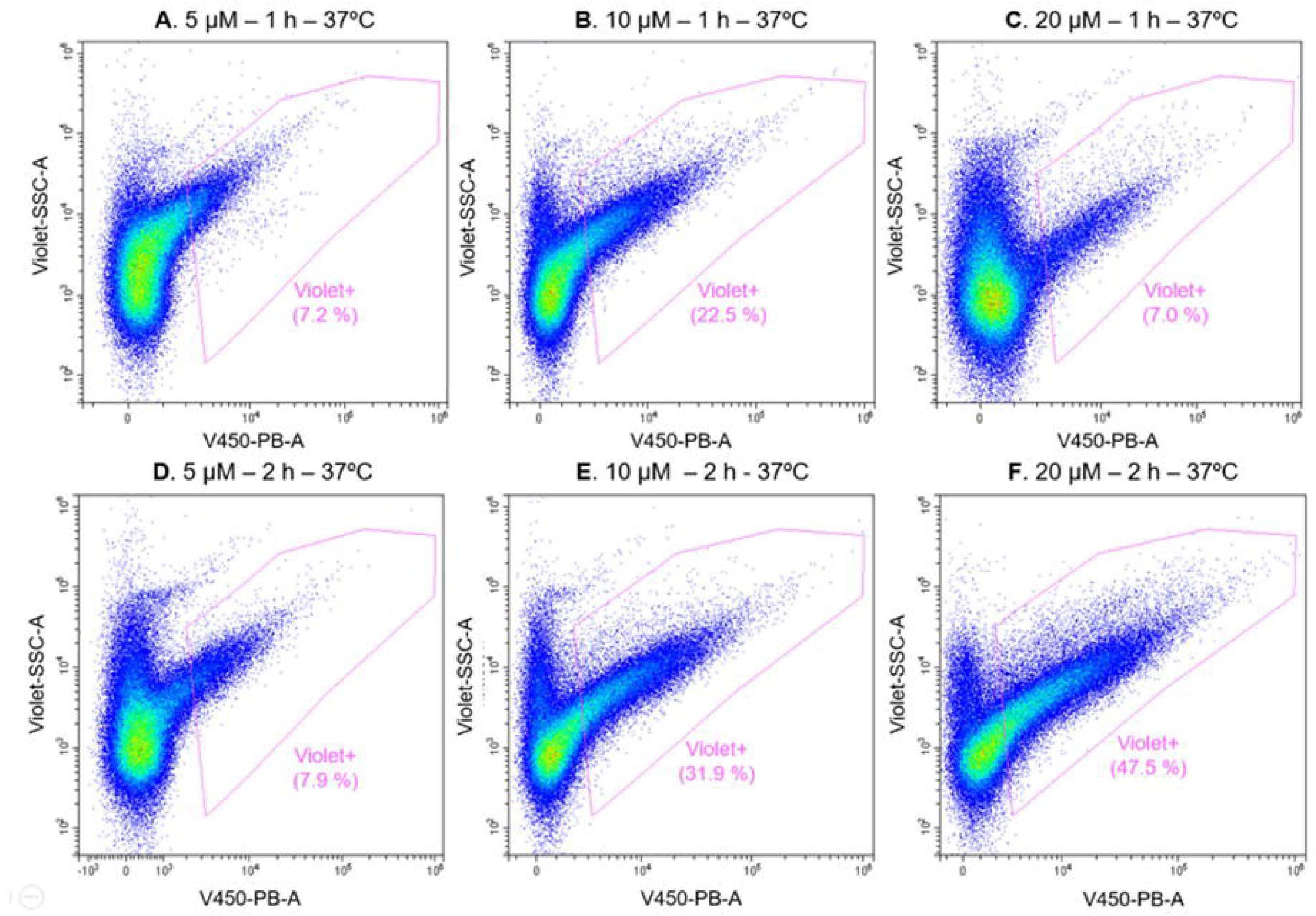

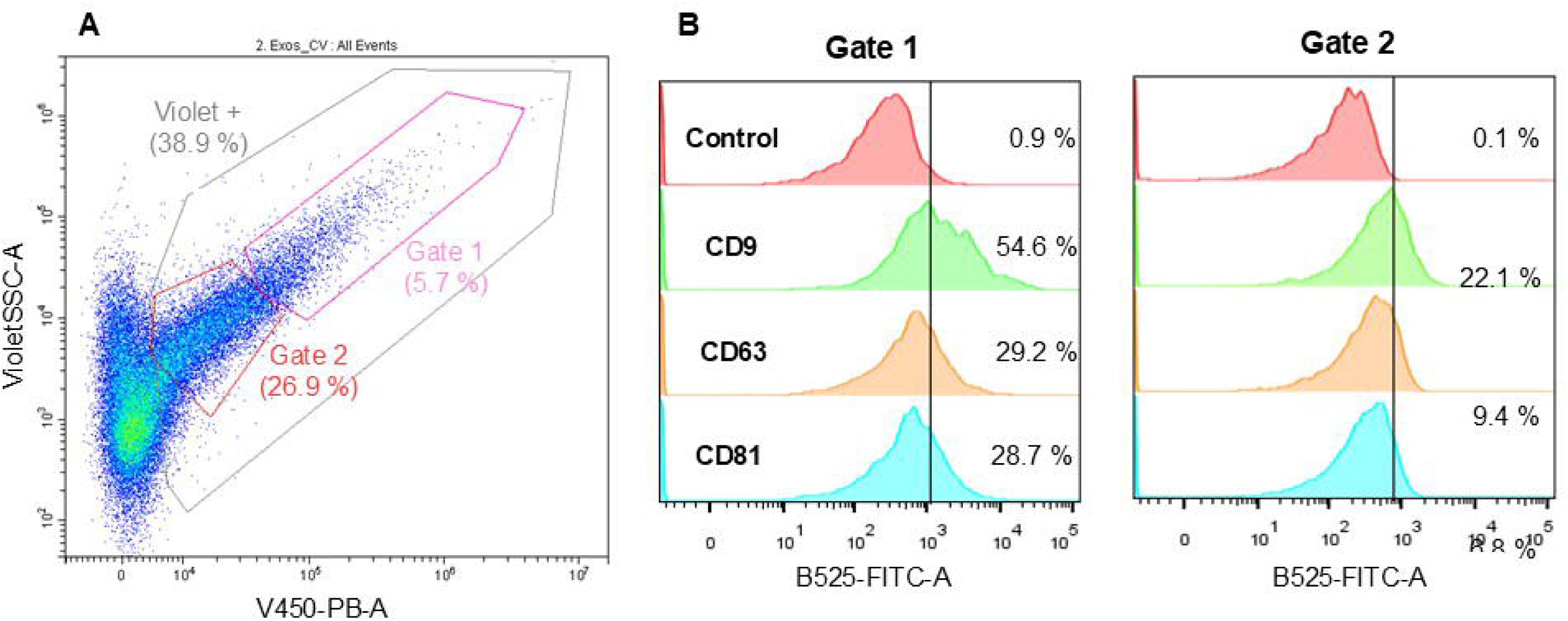

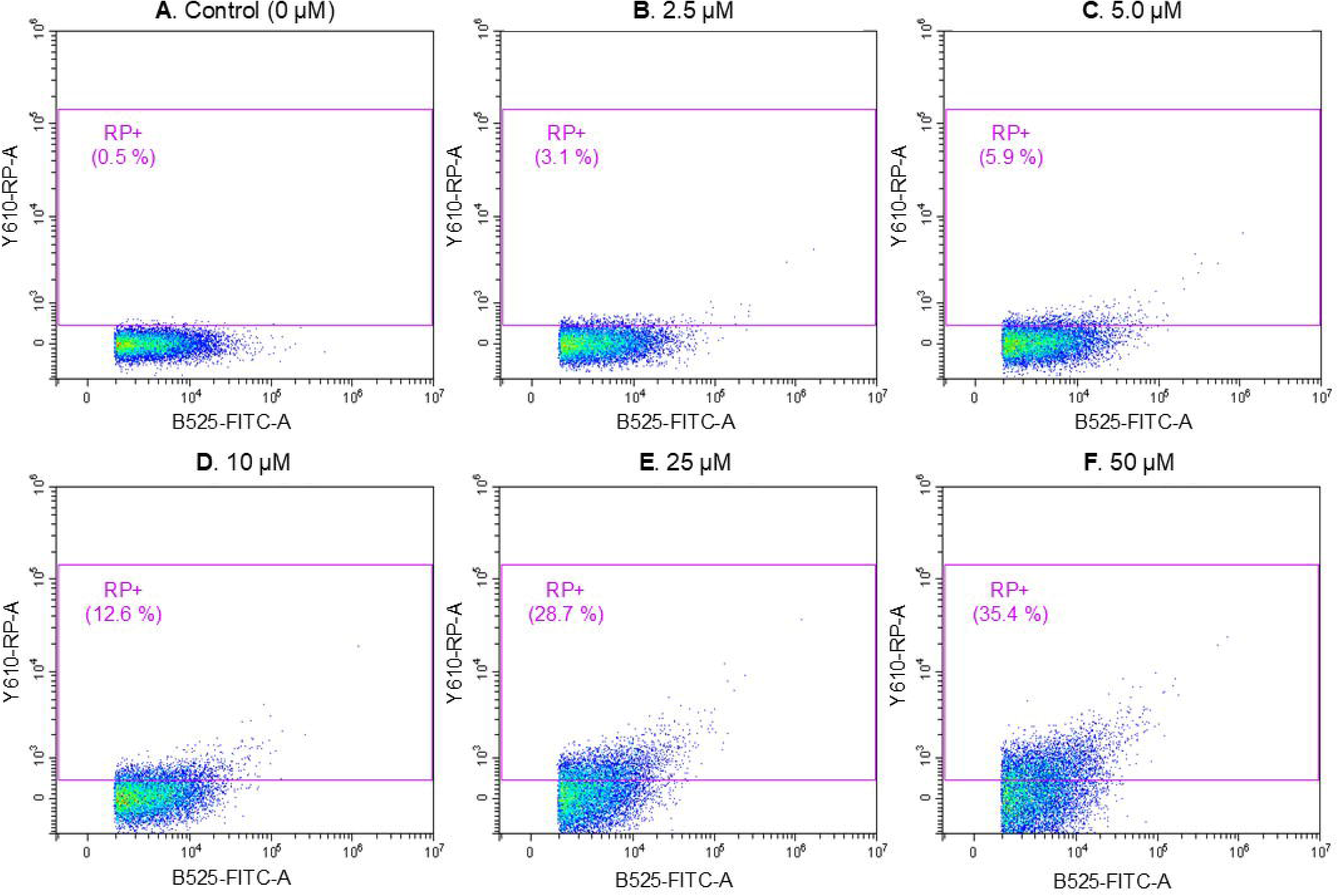

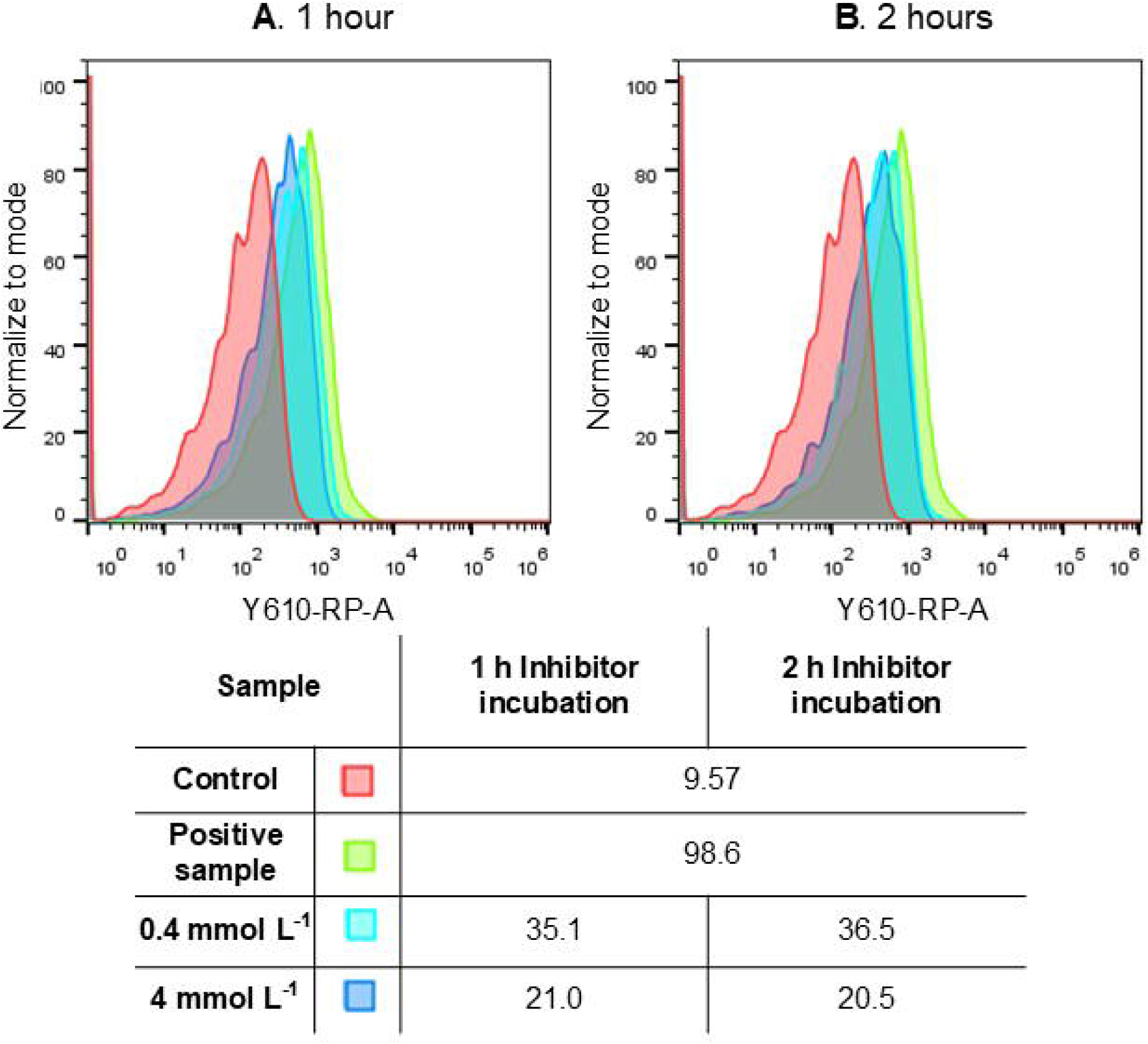

