## Supplemental File for "Aldehyde dehydrogenase 1A3 detection in extracellular vesicles from breast cancer cell lines by nano-flow cytometry"

### Supplementary Data

#### S1. Buffers and solutions

4-(2-hydroxyethyl)-1-piperazineethanesulfonic acid (HEPES, ref. 252859), DL-dithiothreitol (DTT, ref. D0632), sodium phosphate dibasic (ref. 71636), potassium phosphate dibasic (ref. 795496), sodium chloride (ref. S3014), potassium chloride (ref. P3911), magnesium chloride (ref. M8266), boric acid (ref. B6768), glycine (ref. 50046), and bovine serum albumin (BSA, ref. A4503) were purchased from Sigma-Aldrich (Merck KGaA, DE). Sodium carbonate (ref. 131648) was purchased from Panreac (ES). Skimmed milk was purchased from local supplier (Nestlé Sveltesse). All solutions were prepared with ultrapure MilliQ water (Millipore® System, resistivity 18.2 MΩ·cm).

The composition of the buffers and solutions was:

- HEPES buffer: 25 mmol $L^{-1}$ HEPES, 25 mmol $L^{-1}$ $MgCl_2$, pH 7.4.
- HEPES reaction buffer: 50 mmol $L^{-1}$ HEPES, 50 mmol $L^{-1}$ $MgCl_2$, 5mM DTT, pH 8.0.
- PBS 1x buffer: 10 mmol $L^{-1}$ $Na_2HPO_4$, 137 mmol $L^{-1}$ NaCl, 2.7 mmol $L^{-1}$ KCl, 1.8 mmol $L^{-1}$ $K_2HPO_4$, pH 7.4.
- PBS - 0.5 % BSA buffer: 0.5 % w/v BSA in PBS 1x buffer.
- Skimmed milk 3 % in PBS: 3 % w/v of skimmed milk in PBS 1x buffer.
- Carbonate/bicarbonate buffer: 100 mmol $L^{-1}$ $Na_2CO_3$, pH 9.6.
- Boric acid buffer: 100 mmol $L^{-1}$ $H_3BO_3$, pH 8.5.
- Glycine blocking solution: 500 mmol $L^{-1}$ glycine in PBS 1x buffer.

#### S2. Cell culturing, exosome isolation and purification

The cell lines used were breast cancer cell lines SKBR3 (ATCC, ref. HTB-30), MDA-MB-231 (ATCC, ref. HTB-26) and MCF7 (ATCC, ref. HTB-22). Expansion of cell population was carried out from 5 x $10^6$ cells in T-175 flask containing 35 mL of Dulbecco's Modified Eagle's medium. The media were supplemented with 10 % exosome-depleted fetal bovine serum (FBS) and 100 U $mL^{-1}$ penicillin-streptomycin. The temperature was maintained at 37 °C in a humidified, concentrated $CO_2$ (5 %)

atmosphere. Once cells reached approximately 95 % confluence on the T-175 flask, the culture supernatant was removed and stored at –21ºC until to exosome isolation.

Exosomes were purified according as previously reported by our research group.(Moura et al., 2020) The supernatant from the SKBR3, MDA-MB-231, MCF7 and A549 cell lines were subjected to differential centrifugation as follows: 2,000 x *g* for 15 minutes (removal of residual cells) and 10,000 x *g* for 30 minutes (removal of cellular debris, large and medium-sized EVs). Then, a Beckman Coulter Optima L-80XP ultracentrifuge at 100,000 x *g* for 60 minutes, either with a 70Ti or 50.2Ti rotor to pellet exosomes and other small EVs. After that, the supernatant was carefully removed, and crude exosome-containing pellets were resuspended in 1 mL of HEPES buffer (pH 7.4, 0.22 µm filtered and sterile) and pooled. The second round of the same ultracentrifugation setting was carried out, and the resulting exosome pellet resuspended in 250 µL (per each 100 mL of supernatant) of HEPES buffer (pH 7.4, 0.22 µm filtered and sterile) and stored at –21 °C. All centrifugation steps were performed at a temperature of 4 °C.

### S3. Intracellular staining of ALDH in breast cancer cells

To evaluate the presence of ALDH enzyme in breast cancer cells from SKBR3, MDA-MB-231 and MCF7, an intracellular staining was performed as reference method.

Firstly, the fixation of 3 x $10^5$ cells from culture was done by incubation with 200 µL of diluted formaldehyde 4 % in PBS for 10 minutes at room temperature. Then, the microplate was centrifuged (3 min at 800 x *g*), supernatant was discarded, and cells were incubated with 200 µL of PBS containing 2 % of FBS and 0.1 % of saponin, for 10 minutes at room temperature. Then, the supernatant was discarded after centrifugation, and cells were resuspended in FBS containing 0.3 % of saponin for permeabilization of cellular membranes. Then, fixed and permeabilized cells were incubated with 2.5 µg $mL^{-1}$ (3 µL) of specific rabbit polyclonal antibodies against ALDH1A3 for 45 minutes at room temperature. At the same time, intracellular positive control samples were incubated at the same conditions with mouse monoclonal antibodies against vinculin, an ubiquitous cytoskeletal protein of mammalian cells.

Then, two washing steps were performed with PBS containing 2 % of FBS and 0.1 % of saponin, and the cells were resuspended in the same buffer containing 8 µg $mL^{-1}$ (1 µL) of secondary antibodies antirabbit-PE and incubated for 30 minutes at room temperature. In the case of the control samples, they were incubated with antimouse-Cy5 at the same conditions. Finally, two washing steps were performed, and the cells were resuspended in 200 µL PBS 1x buffer and analyzed by flow cytometry.

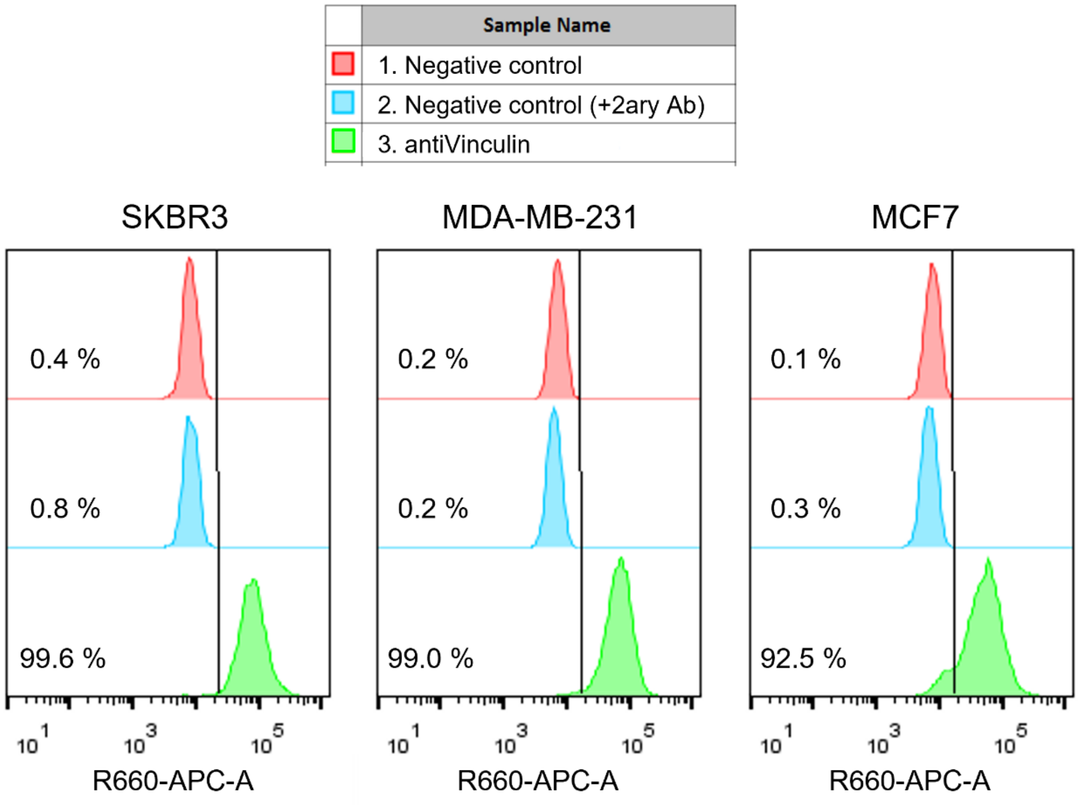


Figure S1: Histograms from intracellular staining of cells from SKBR3, MDA-MB-231 and MCF7 breast cancer cell lines with antiVinculin antibodies, labelled with Cy5-modified antimouse secondary antibodies. In red color, negative controls of the experiment, only cells; in blue color, control with cells only incubated with secondary antibodies; in green color, cells with antiVinculin antibodies plus labelled secondary antibodies.

As a positive control of the permeabilization of cellular membranes, and to confirm their integrity, breast cancer cells were analyzed with antibodies against vinculin. Figure S1 shows histograms of the samples, in all cases with more than 90 % of positivity. In detail, 99.6 % in SKBR3 cells, 99.0 % in MDA-MB-231 cells, and 92.5 % in MCF7 cells.

### S4. Characterization of EVs by nanoparticle tracking analysis, cryogenic transmission electron microscopy and BCA protein assay

The NTA analysis of EV samples derived from breast cancer cell lines SKBR3, MDA-MB-231 and MCF7 revealed clear peaks between 100 and 200 nm, the size expected for exosomes. The size distribution of SKBR3 exosomes only showed a peak at 125 nm (Fig. S2A) of individual vesicles. On the other hand, the size distributions of MDA-MB-231 and MCF7 exosomes size showed similar histograms, with a major peak at 145 nm and 115 nm, respectively, corresponding to individual vesicles. MDA-MB-231 showed two lower peaks at 215 and 325 nm, corresponding to small aggregates (Fig.

S2B). Meanwhile, the size distribution of MCF7 exosomes also showed some aggregates, although at smaller diameters of 155 nm and 205 nm (Fig. S2C). The exosomes from the three breast cancer cell lines were further analyzed by Cryo-TEM, confirming the presence of individual and small aggregates of vesicles in the range between 50 and 400 nm, as shown in Figure S2.

Particle concentration of the samples used for this study was estimated by NTA, and total protein concentration was estimated by BCA protein assay, as shown in Table S1. Exosome sample derived from SKBR3 cells was more concentrated ($4.30 \pm 0.05 \cdot 10^{11}$ particles $mL^{-1}$ and 0.874 mg $mL^{-1}$) than the other two samples. In the case of MDA-MB-231 cells, NTA showed a concentration approximately 8 times lower than that of SKBR3 cells ($5.65 \pm 0.22 \cdot 10^{10}$ particles $mL^{-1}$), being protein concentration only 4 times lower (0.220 mg $mL^{-1}$). On the other hand, in MCF7 exosome samples, both measurements agree, being approximately 4 times lower ($1.64 \pm 0.08 \cdot 10^{11}$ particles $mL^{-1}$, and 0.221 mg $mL^{-1}$). Although NTA is the gold-standard method for exosome counting, it cannot differentiate between vesicles and protein aggregates and may lead to overestimated results.

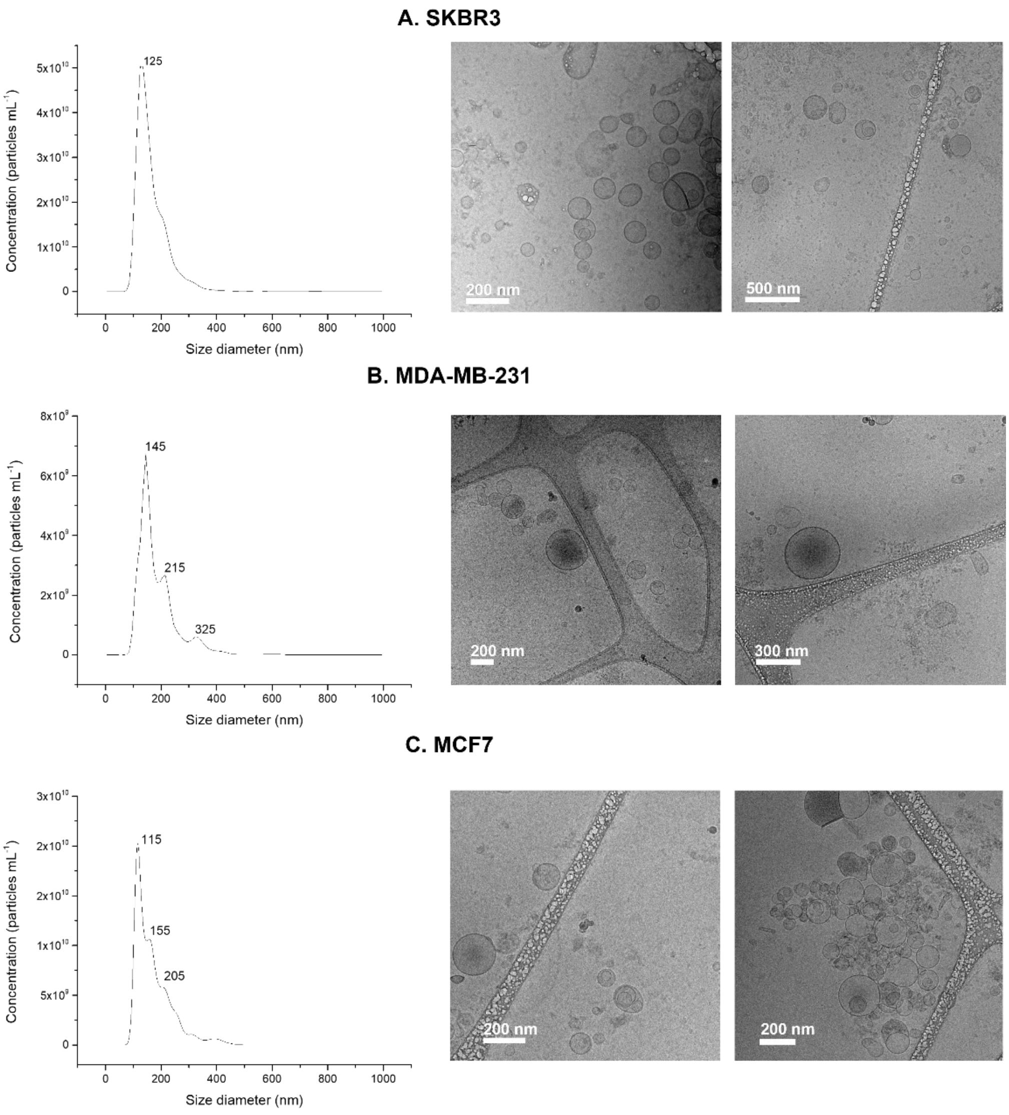


Figure S2. Characterization by NTA and Cryo-TEM micrographs of purified exosome samples from SKBR3 (panel A), MDA-MB-231 (panel B), and MCF7 (panel C) cancer cell lines. The NTA characterization analyzed raw data videos by triplicate during 60 s with 25 frames per second and the temperature of the laser unit set at 24.8°C. Cryo-TEM images were obtained at an acceleration voltage of 200 kV.

Table S1: NTA particle concentration and total protein concentration of EVs samples

| SAMPLE | PARTICLE CONCENTRATION (PARTICLES ML$^{-1}$) | PROTEIN CONCENTRATION (MG ML$^{-1}$) |
|---|---|---|
| **SKBR3** | 4.30 / SD 0.05 $\cdot 10^{11}$ | 0.874 |
| **MDA-MB-231** | 5.65 / SD 0.22 $\cdot 10^{10}$ | 0.220 |
| **MCF7** | 1.64 / SD 0.08 $\cdot 10^{11}$ | 0.221 |

### S5. Characterization of exosomes by bead-based flow cytometry assay

Conventional flow cytometry was used to estimate the presence of general protein markers of exosomes in the membranes of the EVs derived from SKBR3, MDA-

MB-231 and MCF7 breast cancer cell lines. The presence of tetraspanin receptors CD9, CD63 and CD81 was evaluated, following MISEV2018 guidelines(Théry et al., 2018), to confirm the presence of exosomes in the samples. The vesicles were immobilized on magnetic particles as solid support, following previously reported protocols.(Moura et al., 2020)

The immobilization of exosomes on Dynabeads M450 tosylactivated superparamagnetic particles (MPs) were performed as follows: 3.5 x $10^{10}$ exosomes were added to 40 µL of MPs, equivalent to 1.6 x $10^{7}$ particles. The reaction was carried out in 0.1 mol $L^{-1}$ borate buffer pH 8.5, in order to ensure the nucleophilic reaction by the amine group. The incubation step was performed overnight with gentle shaking (~900 rpm) at 4ºC. After that, 0.5 mol $L^{-1}$ glycine solution was added to ensure the blocking of the any remaining tosylactivated groups, by an incubation for 2 h at room temperature. After that, the exosomes-modified magnetic particles (exosomes-MP) were resuspended in 160 µL of 10 mmol $L^{-1}$ PBS buffer to dilute the MPs suspension at 1 x $10^{5}$ MPs per µL. The exosomes-MP were maintained at 4ºC until use.

The presence of the CD9, CD63 and CD81 biomarkers was investigated using FITC-modified primary antibodies. The direct labeling of 5 x $10^{5}$ exosome-modified MPs was performed by incubation with 5 µL (5 µg $mL^{-1}$) of primary antibodies (*i.e.,* FITC-modified antiCD9, antiCD63 and antiCD81), for 30 min with gentle shaking at 4ºC. After that, three washing steps with PBS buffer containing 0.5 % BSA were performed. The labeled MPs were resuspended in 500 µL of PBS 1x buffer and measured by flow cytometry.

The presence of tetraspanin markers on the surface of exosomes derived from SKBR3, MDA-MB-231 and MCF7 cell lines immobilized on MPs was determined by flow cytometry as shown in Figure S3. The presence of CD9, CD63 and CD81 as ubiquitous protein biomarkers was successfully identified in the three samples. FITC-modified mouse monoclonal antiCDX (being CDX CD9, CD63 and CD81) antibodies were used. As a control of specific staining, anti-CDX antibodies were incubated with negative control MPs (protein-modified MPs, without exosomes). This control sample show less than 1 % of positivity for each of the three FITC-modified anti-tetraspanin antibodies (*i.e.,* 0.7 % CD9, 0.3 % CD63, and 1.0 % CD81).

The results from the bead-based assay showed positive signals in the three exosomes samples with the three markers investigated. Specifically, exosomes derived from SKBR3 show 100 % of positivity in CD9, 51 % in CD63, and 44 % in CD81 markers. Next, exosomes derived from MDA-MB-231 show 39 % of positivity in CD9, 64 % of

CD63, and 38 % in CD81. And finally, exosomes derived from MCF7 show a 92 % of positivity in CD9, 67 % in CD63 and 48 % in CD81. The results confirm the presence of these biomarkers on the exosomes membranes.

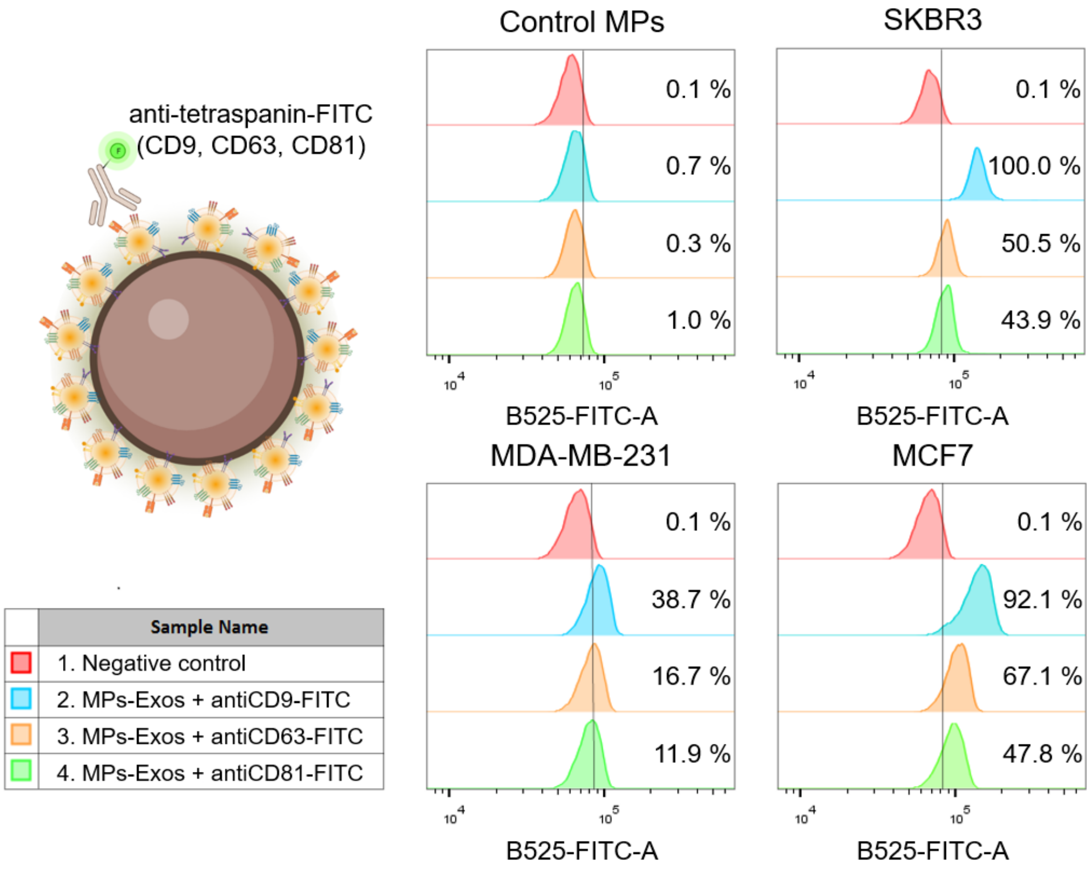


Figure S3: Bead-based flow cytometry assay for the characterization of protein surface markers in exosomes derived from breast cancer cell lines. Magnetic particles were covalently modified with exosomes derived from SKBR3, MDA-MB-231, and MCF7. Specific antibodies against tetraspanins CD9, CD63 and CD81, modified with FITC were used. As control sample, antibody modified magnetic particles, blocked with glycine, were used to verify the specificity of the immunoaffinity reaction.

### S6. Sandwich ELISA for the determination of ALDH

Exosomes derived from SKBR3 breast cancer cell line were subjected to sandwich ELISA determination of ALDH1A3 presence on their membranes. Besides, the integrity of the vesicles and well-functioning of the ELISA method was evaluated by reacting with antiCD81 as general exosome membrane biomarker.

As shown in section 4.2 with intracellular staining and flow cytometry, SKBR3 cells displayed a high expression of ALDH1A3 in the cytosol. To evaluate the presence of the ALDH1A3 enzyme in the membranes of SKBR3-derived exosomes, a sandwich ELISA was used as a reference and standard method for determining protein markers.

As expected, the sandwich ELISA using antiALDH1A3 as capture antibody did not show higher signals in the presence of SKBR3 exosomes, as shown in Figure S4. Meanwhile, a calibration curve was determined with the same exosomes samples captured by antiCD81 antibodies and detected by antiCD63-HRP, at the same experimental conditions. In conclusion, the results confirm that exosomes derived from SKBR3 cell line did not express ALDH1A3 in their membranes. Therefore, the exosomes show a similar ALDH compartmentalization as the cells, that locate the enzyme only in the intracellular space, in agreement with previous studies. (Zanoni et al., 2022)

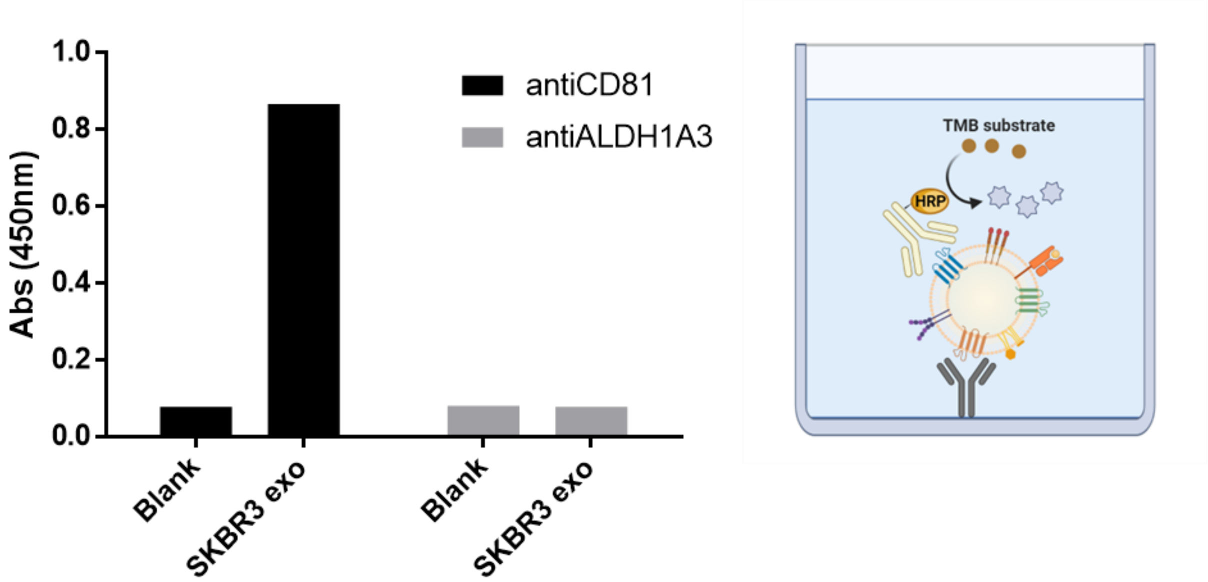


Figure S4: Sandwich ELISA determination of ALDH1A3 presence in the membranes of exosomes derived from SKBR3 breast cancer cell line. Antibodies against ALDH1A3 were used to capture the exosomes, further labelled by antiCD63-HRP as ubiquitous membranes protein marker.

Firstly, specific antibodies against ALDH1A3 (1:250 dilution factor, 0.4 µg $mL^{-1}$) and antiCD81 (1:250 dilution factor, 2 µg $mL^{-1}$) were diluted in carbonate/bicarbonate buffer and were incubated (100 µL per well) overnight at 4ºC in Maxisorp polystyrene microplates. Then, the coating solution was removed, and remaining sites were blocked (skimmed milk 3 % in PBS) for 2 hours at room temperature and gentle shaking (550 rpm). Blocking solution was removed and plates were washed with PBS – 0.5 % BSA three times. Meanwhile, six different exosomes samples were prepared in PBS, with concentrations ranging from $3.42 \cdot 10^8$ to $1.07 \cdot 10^7$ particles $mL^{-1}$, according to NTA measurements. The samples were incubated by triplicates for 1 hour at room temperature and gentle shaking. Then, supernatants were carefully removed, and three washing steps were applied. To label the vesicles, HRP-conjugated antiCD63 (1:1000 dilution factor, 0.3 µg $mL^{-1}$) antibodies were incubated for 1 hour at room temperature and gentle shaking. Again, supernatants were carefully removed, and three washing steps were applied. Finally, TMB substrate solution (1:1 TMB and $H_2O_2$ from Pierce TMB substrate kit) was incubated for 30 minutes at room temperature. The reaction was stopped by adding 2 mol $L^{-1}$ sulfuric acid and the colored product was quantified spectrophotometrically at 450 nm.

### S7. Nano-Flow cytometry studies of breast cancer exosomes

#### *S7.1 CFSE and Violet staining of exosomes*

For this study, CellTrace™ CSFE proliferation kit (excitation 488 nm, emission 525 nm, ref. C34554, Thermo Fisher Scientific) and CellTrace™ Violet proliferation kit (excitation 425 nm, emission 450 nm, ref. C34557, Thermo Fisher Scientific) were used for exosome staining (Fig. S5). Exosomes derived from SKBR3 breast cancer cell line were used to optimize several experimental parameters, including the concentration, the temperature and time of incubation and some cytometer parameters such as threshold and gains. For the CFSE assay, exosomes were incubated with CFSE 20 µmol $L^{-1}$ for 30 minutes, 1 hour and 2 hours at 37ºC to study the incubation time, following previous studies.(Ender et al., 2020; Hoen et al., 2012) Additionally, a sample with exosomes and CFSE 20 µmol $L^{-1}$ was incubated for 2 hours at RT to study the incubation temperature. All tubes were diluted to 1 mL with HEPES buffer and measured by nano-flow cytometry. Regarding the instrumental parameters, different channel gains and thresholds were analyzed with exosomes samples. As shown in Figure S5, the best discriminated exosome population when comparing all the previously mentioned parameters was the sample incubated with CFSE for 2 hours at 37ºC and a B525 (FITC) channel gain of 1,000 units (Fig S6, panel E).

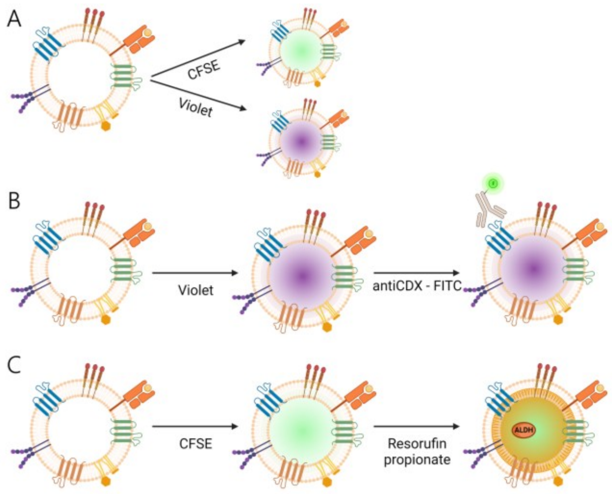


Figure S5: Scheme of the exosome staining methods studied. On Panel A, to identify the exosomes on nano-flow cytometry experiments, the vesicles were labelled with protein-binding fluorescent reporters as CFSE or Violet tracers. On Panel B, to determine the presence of membrane protein biomarkers, Violet-labelled exosomes were incubated with FITC-conjugated primary antibodies. On Panel C, to assess the intrinsic activity of ALDH enzymes inside the vesicles, CFSE- exosomes were incubated with resorufin propionate substrate.

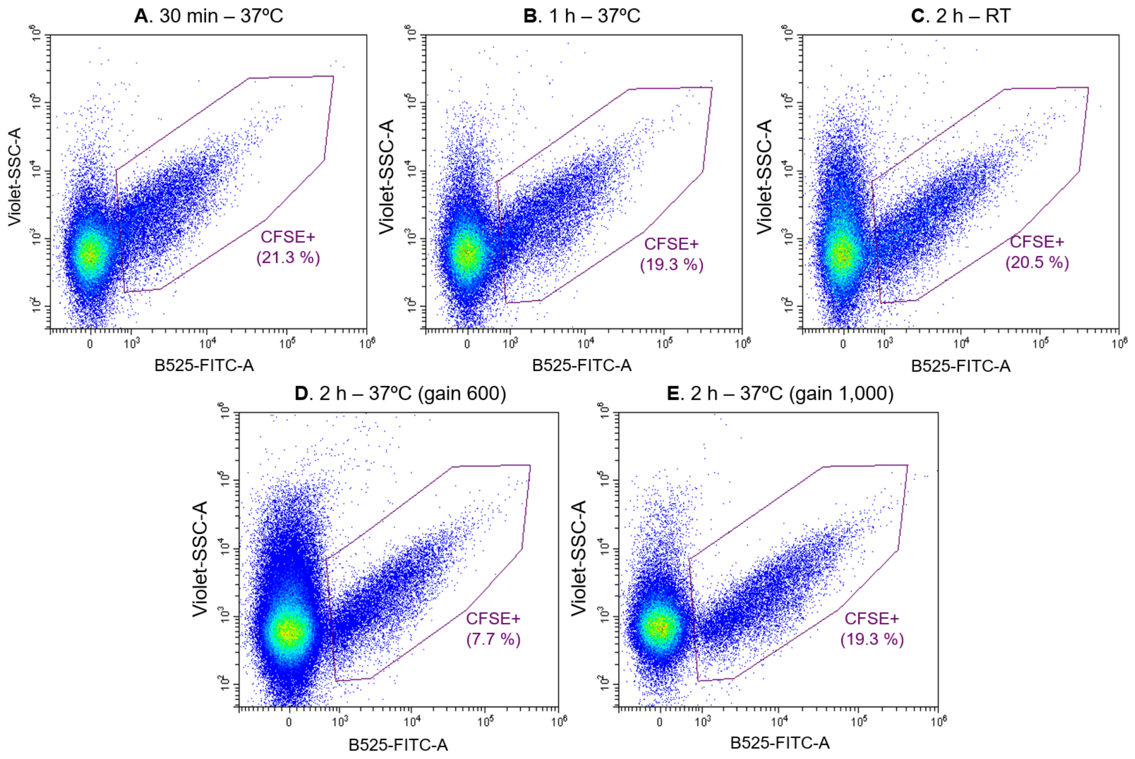


Figure S6: Optimization of the CFSE labelling of SKBR3-derived exosomes. Dot plot representations (on x axis: B525-FITC-A channel signal, on y axis Violet-SSC-A). Incubations with CFSE were performed at 37ºC for 30 minutes (Panel A), 1 hour (Panel B), 2 hours (Panel D and E). On Panel C, the incubation was done for 2 hours at room temperature. The gain of B525 channel was optimized to 1,000 units (Panel E).

In parallel, serial dilutions of SKBR3 exosomes incubated with 20 µmol $L^{-1}$ of CFSE for 2 hours at 37ºC were measured to test the exosomes concentration range suitable for nano-flow cytometric measurements. As shown in Figure S7, the concentration of the sample clearly affects the abort rate of the measurements, directly related with the quality of the results. Briefly, if two particles enter the laser beam too close together, and the instrument cannot detect them independently, the instrument will abort them both. Following Cytoflex LX manufacturer technical recommendations, an abort rate below 8.0 % is considered as good quality results in the case of nanoparticles samples. In our case, the best results were obtained with a 100-fold dilution of the exosomes sample, shown in Figure S7, panel C. The NTA count of SKBR3 exosomes sample allows to calculate the corresponding concentration of particles in the measured tube, which was $6.45 \cdot 10^{7}$ particles $mL^{-1}$.

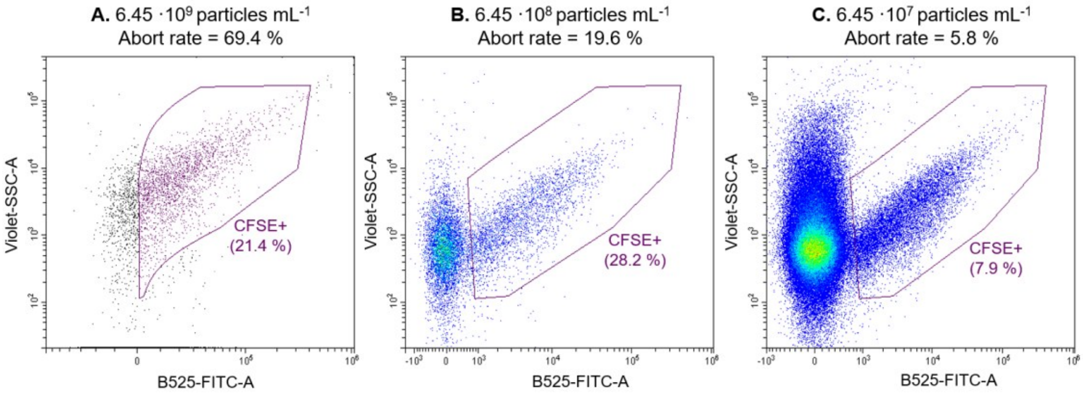


Figure S7: Optimization of SKBR3-derived exosomes concentration on nano-flow cytometry measurements. Dot plot representations (on x axis: B525-FITC-A channel signal, on y axis Violet-SSC-A). 10-fold serial dilutions of SKBR3 exosomes CFSE labelled. The concentrations were calculated according to NTA particle counting.

To further test the CFSE labelling protocol, as well as the exosomes concentration range suitable for the nano-flow cytometry measurements, exosomes derived from various cell lines were measured, as shown in Figure S8. As can be observed, variable percentages of CFSE+ vesicles were obtained. This fact can be related with the exosomes concentration, as more diluted samples had to be recorded for longer times, therefore, increasing the noise signals measured. As previously explained, the acquisition settings of the cytometer were configured to measure a minimum of 10,000 particles in the CFSE+ gate.

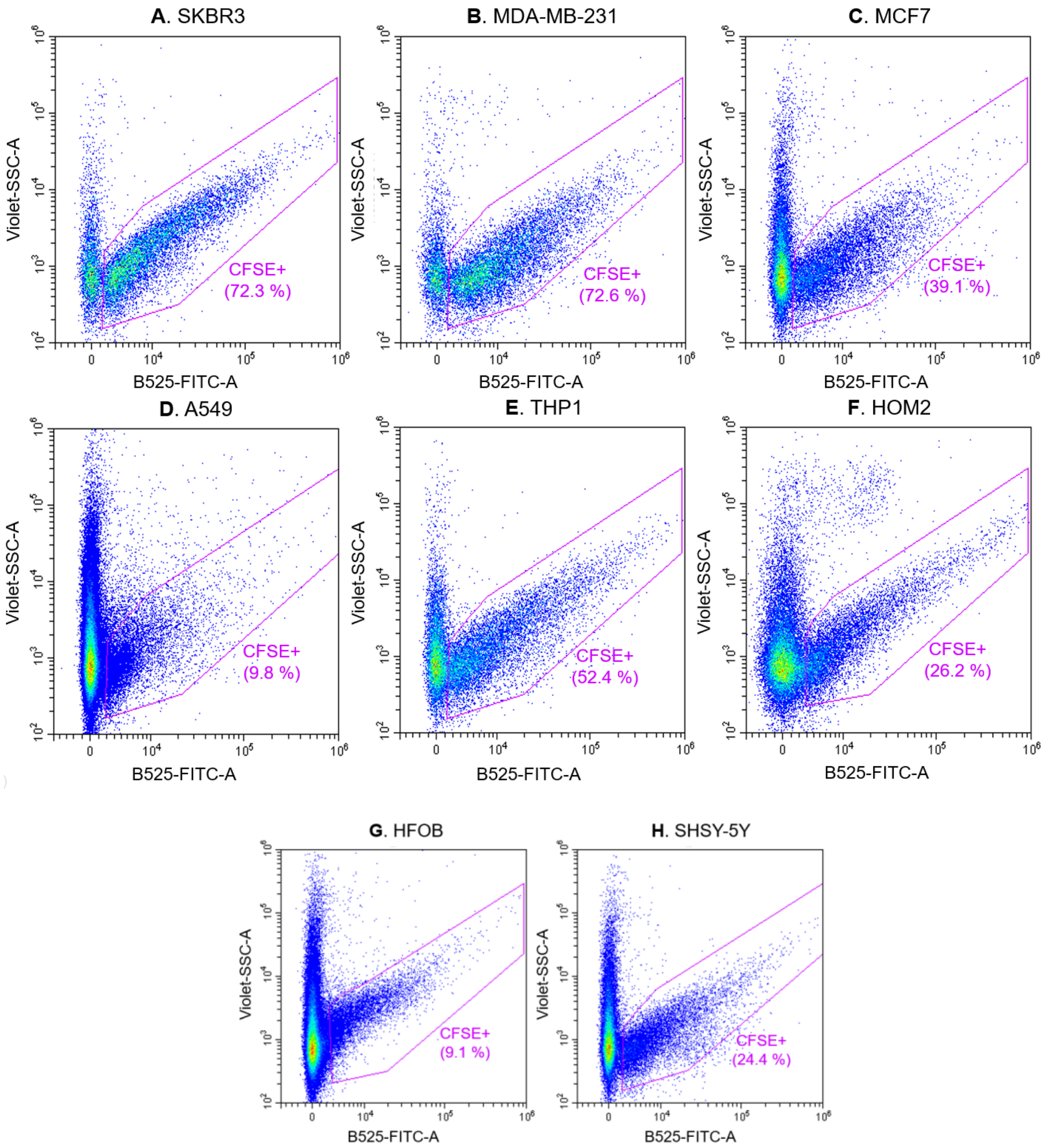


Figure S8: Nano-flow cytometry measurements of exosomes derived from cell culture supernatants. Dot plot representations (on x axis: B525-FITC-A channel signal, on y axis Violet-SSC-A). CFSE labelled exosomes derived from SKBR3 (Panel A), MDA-MB-231 (Panel B) and MCF7 (Panel C) breast cancer cell lines; from A549 (Panel D) lung cancer cell line; from THP1 (Panel E) monocytic cell line; from HOM2 (Panel F) lymphoblastoid cell line; from HFOB (Panel G) osteoblasts cell line; and SHSY-5Y (Panel H) neuroblastoma cell line.

A comparison was done between the particle counting by Nanoparticle tracking analysis and CFSE labelled particles in nano-flow cytometry (concentration of CFSE+ particles subpopulation), as shown in Figure S9 bar graph and table. It can be observed that there is not a consistent labelling ratio between the cell lines. The differences might be related to different methodological issues, as the overestimation of particle concentration due to vesicles and protein aggregates in NTA.

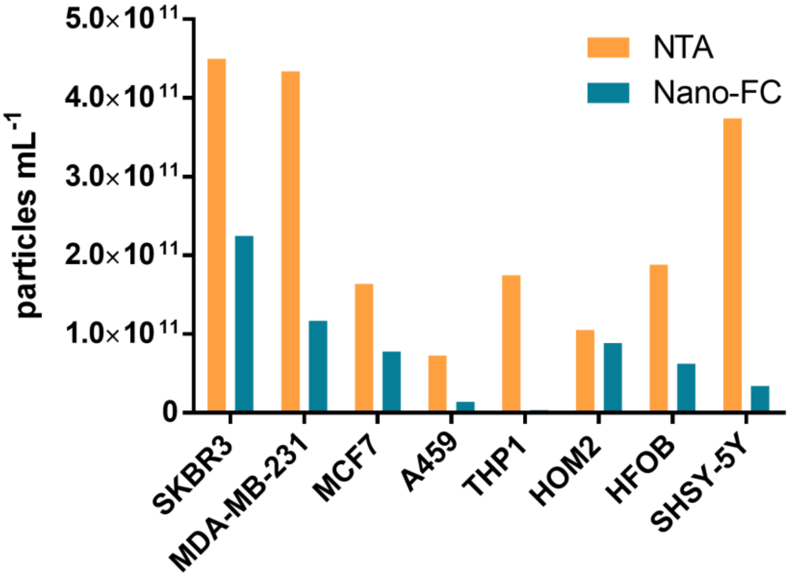


| Exosomes cell line | NTA (particles $mL^{-1}$) | Nano-FC (particles $mL^{-1}$) | % Nano-FC vs. NTA |
|---|---|---|---|
| **SKBR3** | 4.50e+11 / SD 3.13e+09 | 2,25E+11 | 50,0% |
| **MDA-MB-231** | 4.34e+11 / SD 7.75e+09 | 1,17E+11 | 27,0% |
| **MCF7** | 1.64e+11 / SD 7.55e+09 | 7,76E+10 | 47,3% |
| **A459** | 7.25e+10 / SD 2.69e+09 | 1,39E+10 | 19,2% |
| **THP1** | 1.75e+11 / SD 7.14e+09 | 3,64E+09 | 2,1% |
| **HOM2** | 1.05e+11 / SD 3.87e+09 | 8,87E+10 | 84,5% |
| **HFOB** | 1.88e+11 / SD 5.16e+09 | 6,24E+10 | 33% |
| **SHSY-5Y** | 3.74e+11 / SD 1.68e+10 | 3,39E+10 | 9,1% |

Figure S9: Comparison of NTA vs Nano-flow cytometry. Both bar graph and table show the particle concentration determined by each method of eight different cell lines: SKBR3, MDA-MB-231, and MCF7 breast cancer cell lines; A549 lung cell line, THP1 monocyte cell line, HOM2 lymphoid cell line, HFOB osteoblast cell lines, and SHSY-5Y neuroblastoma cell line. The ratio of particle concentration determined by nano-flow cytometry versus NTA.

In the case of Violet staining, exosomes derived from SKBR3 were incubated with CellTrace Violet at concentrations of 5 µmol $L^{-1}$, 10 µmol $L^{-1}$ and 20 µmol $L^{-1}$ for 1 hour and 2 hours at 37ºC. With the knowledge of CFSE staining optimization results, all incubation were performed at 37ºC. All tubes were diluted to 1 mL with HEPES buffer and measured by nano-flow cytometry. As can be seen in Figure S10 when all the previously mentioned parameters were compared, the incubation of the exosomes for 2 hours at 37ºC in the presence of Violet staining at a concentration of 20 µmol $L^{-1}$ was the best condition to discriminate the exosomes from the background noise, as determined for CFSE (47.5 %). The gain of the V450 (Violet) channel was optimized manually to 500 units (Fig. S10).

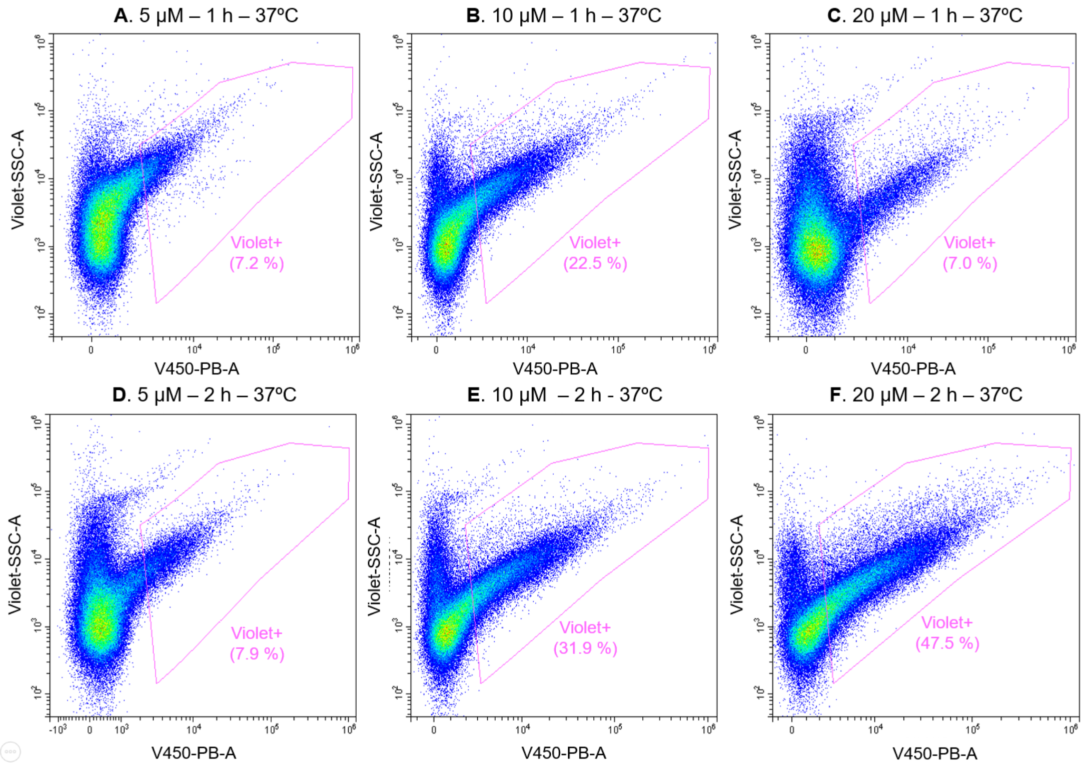


Figure S10: Optimization of the Violet labelling of exosomes. Dot plot representations (on x axis: V450-PB-A channel signal, on y axis Violet-SSC-A channel). Incubations with Violet at 5, 10 and 20 µmol $L^{-1}$ staining were performed at 37 ºC for 1 hour (Panel A, B, C) and 2 hours (Panel D, E, F).

### *S7.2 Experimental design of ALDH determination of tracer-labelled exosomes by nano-flow cytometry*

To do that, exosomes derived from SKBR3 were stained with CellTrace violet reagent and afterwards they were labelled with FITC-modified primary antibodies against CD9, CD63 and CD81. In Figure S11A, a dot plot representing the profile of the exosome sample is shown, using V450-Violet and Violet-SSC-A channels. The distribution of the dots allowed to identify two subgroups selected by the depicted gates in the dot plot graph. The FITC fluorescence of the violet-labelled vesicles, which was due to the labelled antibodies, was then analyzed using B525-FITC channel. As shown in Figure S11B, vesicles selected by Gate 1 were enriched by tetraspanin-expressing vesicles compared to Gate 2, suggesting that Gate1 was selecting vesicles with features of exosomes that is high expression of the tetraspanin, the exosomes canonical markers. As expected from previous results (section S5), CD9 was the most abundant tetraspanin, followed by CD63 and CD81, respectively. For following experiments, the exosomes subpopulation was selected according to 'Gate 1'.

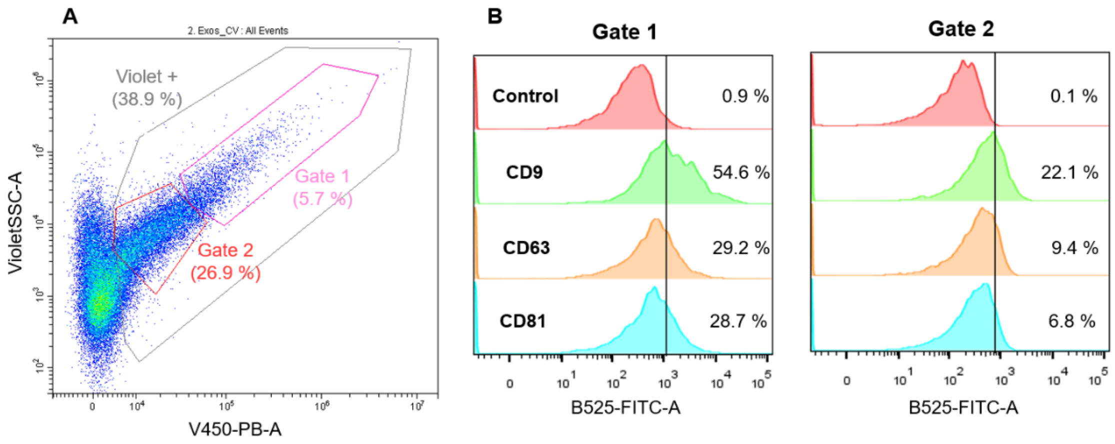


Figure S11. Analysis of the membrane protein markers of exosomes derived from SKBR3 breast cancer cell line. Panel A shows a dot plot representation of a measurement with the different gates: in grey, the total vesicles violet labelled; in pink, Gate 1 subpopulation; and in red, Gate 2 subpopulation. Panel B and C show the tetraspanin expression of both subpopulations, respectively, determined by B525-FITC-A fluorescent signals.

With the nano-flow cytometry, the tracer labelling combined with the specific-antibody staining for the exosomes revealed the heterogeneity of the composition of the sample under study. The differences between extracellular vesicles could be related to the different biogenesis pathways that result in the expression of molecular patterns that allow them to be differentiated.

For ALDH activity assays, resorufin propionate (RP) substrate and ALDH specific inhibitor concentrations and incubation times were optimized. In all experiments, all samples were filled until 1 mL with HEPES buffer after incubations and measured by nano-flow cytometry.

Exosomes derived from SKBR3 breast cancer cell lines were incubated with 20 µmol $L^{-1}$ CFSE for 2 hours at 25ºC in the presence of different concentrations of the resorufin propionate substrate, ranging from 2.5 µmol $L^{-1}$ to 50 µmol $L^{-1}$. As shown in Figure S12, the optimal concentration was considered 25 µmol $L^{-1}$ because high intensity signals were observed in Y610-RP channel. The 50 µmol $L^{-1}$ concentration was discarted, considering the potential harmful effect of DMSO (used as diluent in the RP stock solution) to the exosomes.

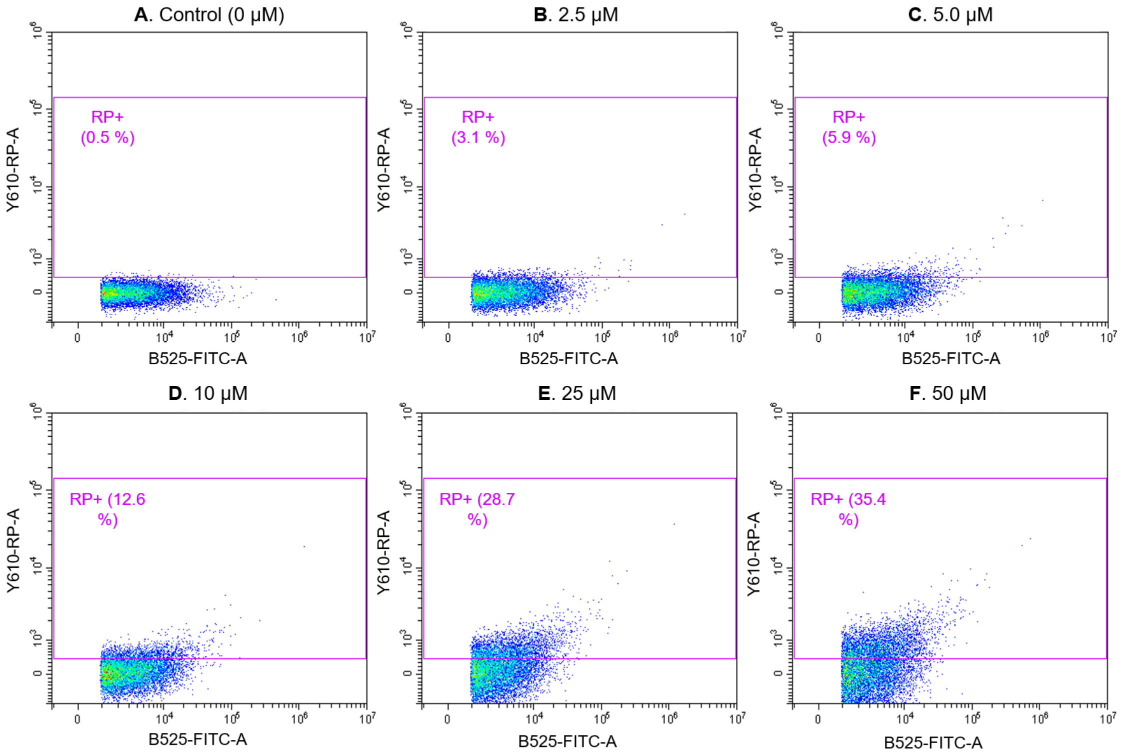


Figure S12: Optimization of resorufin propionate concentration. Dot plot representations (on x axis: B525-FITC-A channel signal, on y axis Y610-RP-A channel). Incubations with different concentrations of resorufin propionate, ranging from 2.5 µmol $L^{-1}$ to 50 µmol $L^{-1}$, plus a negative control sample.

In parallel, the RP incubation time was examined. Both CFSE and resorufin propionate can be simultaneously analyzed by flow cytometry, as their emission spectra are sufficiently distinct to avoid significant overlap. CFSE emits at approximately 517 nm, while resorufin propionate emits around 585 nm. This spectral separation allows for accurate detection in conventional FITC and PE channels, respectively, minimizing the need for extensive compensation. CFSE-labelled SKBR3 exosomes were incubated with 25 µmol $L^{-1}$ RP substrate for different times (30 minutes, 1 hour and 2 hours) at 25ºC (data not shown). The results showed that different incubation times do not increase significantly the resorufin fluorescent signals. Although, an incubation time of 2 hours was chosen to amplify the signal of the enzymatic reaction.

For the ALDH inhibitor assay, SKBR3 exosomes were incubated with CFSE staining and ABD0305 as ALDH specific inhibitor at concentrations of 0.4 mmol $L^{-1}$ (100-fold dilution) and 4 mmol $L^{-1}$ (10-fold dilution) for 1 hour and 2 hours at 37ºC. Then, each sample was incubated with 25 µmol $L^{-1}$ RP for 2 hours. As can be observed in Figure S13, the 4 mmol $L^{-1}$ dilution inhibits more the signal, although it might be also attributed to the effect of ethanol in the biological activity of exosomes. Therefore, in order to prevent unspecific inhibition a concentration of 0.4 mmol $L^{-1}$ was used as working solution with 2 hours incubation time.

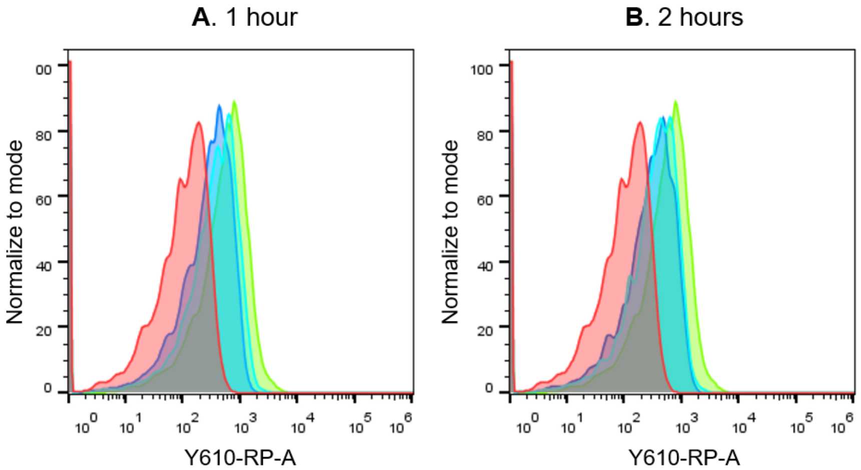


| Sample | | 1 h Inhibitor incubation | 2 h Inhibitor incubation |
|---|---|---|---|
| Control | | 9.57 | |
| Positive sample | | 98.6 | |
| 0.4 mmol $L^{-1}$ | | 35.1 | 36.5 |
| 4 mmol $L^{-1}$ | | 21.0 | 20.5 |

Figure S13: Optimization of ALDH inhibitor incubation. Panel A and B, histogram representations (on x axis: Y610-RP-A channel signal, on y axis: normalized count to mode). On the table, geometric mean of the samples previously represented, with 1 hour inhibitor incubation (Panel A) and 2 hours (Panel B).

Finally, for the comparison of frozen exosomes (stored -21ºC) with non-frozen exosomes (stored at +4ºC), 15 µL of each exosomes sample (SKBR3, MDA-MB-231 and MCF7 cell lines) were incubated with 25 µmol $L^{-1}$ of CFSE and 0.4 mmol $L^{-1}$ ALDH inhibitor for 2 hours at 37ºC. Then, the samples were incubated with 25 µmol $L^{-1}$ of resorufin propionate substrate for 2 hours at 25ºC. Tubes without CFSE, ALDH substrate and ALDH inhibitor were filled with the same correspondent volume with HEPES to have the same conditions.
